# A blastema in sea star larvae integrates wound signaling to drive regeneration-specific and developmental gene expression patterns

**DOI:** 10.64898/2026.09.20.752982

**Authors:** Jon Lee Andrade, Andrew Wolff, Lexi Rios, Adel Fergatova, Veronica Hinman

## Abstract

Whether regeneration depends on the reactivation of developmental programs, regeneration-specific regulatory mechanisms, or both remains a central question in regeneration biology. Here, we investigate these processes in regenerating larvae of the sea star Patiria miniata, a deuterostome with robust regenerative capacity. By integrating single-nucleus transcriptomics with chromatin accessibility profiling across development and regeneration, we identify a regeneration-induced blastema cell state that is molecularly distinct from pre-existing larval populations and serves as the source of regenerated tissues.

We show that regeneration is associated with distinct classes of regeneration-responsive enhancers, including regeneration-specific elements and enhancers reused from development, which link wounding signals to gene regulatory network (GRN) activation. These enhancer classes converge on regulatory programs associated with the transcription factor Runx, positioning Runx as a central node within the inferred regeneration GRN. Notably, we identify a Runx-associated regulatory framework that provides a mechanistic explanation for the de novo emergence of sox4⁺ cells during regeneration through novel deployment of developmentally shared enhancers. Together, our results provide a framework for how wound-induced signals specify regenerative cell states and how regeneration-specific and developmental gene regulatory networks may be coordinated to rebuild lost tissues.

**Teaser:** Distinct enhancer classes link injury signals to developmental gene networks during regeneration

## Introduction

Regeneration is the process by which an organism restores cells, tissues, organs, and whole bodies after acute trauma or injury (Poss & Tanaka, 2024).

This process is widespread among multicellular organisms, but it is unevenly distributed both among and within clades (Bely & Nyberg, 2010; Brockes & Kumar, 2008). Furthermore, the extent to which organisms can regenerate certain structures and the mechanisms by which they do so are highly variable and often context-dependent. Despite this variability, established and emerging systems have been instructive for understanding regeneration at broad scales, though more work is needed to address questions critical to the field.

Two central questions in regeneration biology are how wound-induced signals direct the deployment and specification of regenerative cells, and whether regeneration is driven by regeneration-specific mechanisms that reset cellular states or by the redeployment of developmental pathways to rebuild missing tissues. These questions are particularly tractable in systems with robust regenerative capacity and accessible developmental contexts, such as echinoderms. Indeed, echinoderms have remarkable regenerative potential, with regeneration observed across all classes (Ben Khadra et al., 2018; Carnevali, 2006; Dupont & Thorndyke, 2007; Quispe-Parra et al., 2021; Ferrario et al., 2020). Their regenerative abilities include the replacement of appendages and organ systems, as well as whole-body regeneration (WBR) from detached body fragments, which is relatively rare even among regenerating animals. As deuterostomes, echinoderms occupy a critical position in the evolutionary tree, which could further our understanding of how regeneration evolved in vertebrates, which show weak regenerative abilities compared with some invertebrate groups. Thanks to these characteristics as well as recent improvements in genomic resources and molecular techniques, echinoderms are in an excellent position to rapidly improve our understanding of regeneration (Arshinoff et al., 2022; Cary et al., 2018; Foley et al., 2021; Telmer et al., 2024; Medina-Feliciano & García-Arrarás, 2021).

Echinoderms also possess remarkable regenerative capabilities, including WBR, in their larval stages, which have been observed in sea stars and sea urchins (Kasahara et al., 2019; M. C. Vickery et al., 2001; M. S. Vickery et al., 2002; Wolff & Hinman, 2021). The larva of the bat star, *Patiria miniata* (*P. min*), has in the last decade become an attractive model system for regeneration, providing us with insights into the cellular sources of regenerating tissues, the role of signaling pathways in injury response and axial respecification, and the redeployment of developmental pathways. Bisection of larval *P. min* through the anteroposterior (AP) axis results in two fragments that are both able to regenerate over the course of roughly two weeks, restoring form and function of all missing tissues (Cary et al., 2019). Around 3 days-post-bisection (dpb), cells at the wound site undergo a rapid increase in proliferation coincident with a decrease in proliferation distal to the injury and the expression of various developmentally relevant transcription factors including but not limited to *runt* (now *runx*), *sox2*, and *sox4* (Cary et al., 2019; M. Zheng et al., 2022). This pattern is observed in several regenerative animals in a structure known as the blastema, a wound-proximal collection of proliferative cells that go on to form the new tissues (Can Aztekin, 2024; Ricci & Srivastava, 2018). However, it is unclear whether these cells actually contribute to regenerated tissues in *P. min* and can therefore be considered a blastema, and the lack of a comprehensive, reliable set of molecular markers precludes direct investigation of this population.

Evidence has also supported at least a partial reactivation of embryonic processes at the wound site of larval *P. min*. For example, respecification of the AP axis appears to use embryonic mechanisms as indicated by the upregulation of posterior specification domain transcripts, *wnt8* (now *wnt8a*) and *frizz9/10* (*now fzd9*), in anterior fragments and anterior domain genes, *frizz5/8* (*now fzd5*) and *foxq2* (*now foxe3l*), in posterior fragments (Cary et al., 2019). Furthermore, previous work supports the redeployment of embryonic pathways to respecify serotonergic neurons in regenerating larval *P. min* fragments. This was determined by the sequential expression of the ectodermal stem cell marker, *sox2*, and the neural stem cell marker, *sox4*, followed by *lhx2* (now *lhx9l*), an *elav* homolog, and the production of serotonin, following the progression of the serotonergic neurogenesis pathway in embryos (Cheatle Jarvela et al., 2016; M. Zheng et al., 2022). While this work provides evidence for the reuse of developmental mechanisms, it remains unclear how these embryonic patterns are regulated downstream of wounding and whether any gene regulatory mechanisms are also shared with embryogenesis.

In this study, we demonstrate that wounding induces the formation of a blastema cell population that is molecularly distinct from any cell population present in larvae prior to bisection. We show that this population serves as the source of newly formed tissues during regeneration. In addition, we identify a previously undescribed larval echinoderm cell state that disappears following bisection, although it remains unclear how these cells contribute to regeneration. We further show that the transcriptional program of the blastema is regulated by a combination of regeneration-specific and developmental enhancers. Notably, we find that *sox4* expression is associated with enhancers shared with embryogenesis, and that Runx plays a key role in both the activity of these enhancers and the coordination of regeneration-specific and developmental regulatory mechanisms. Based on these findings, we construct a preliminary gene regulatory network (GRN) that links wounding to the activation of regeneration-specific and developmentally redeployed genes. Together, this work provides a mechanistic framework for understanding how wound-induced signals direct regenerative cell specification and how regeneration-specific and developmental GRNs are coordinated during tissue reconstruction.

## Results

### Cell state atlas of intact *P. min* bipinnaria larvae

As a starting point for understanding how cell states change following bisection and how this might be regulated, we first generated a single-nucleus atlas for intact 9 days-post-fertilization (dpf) bipinnaria larvae using single-nucleus RNA-sequencing (snRNA-seq). Sequencing reads were mapped to the Pmin_3.0 *Patiria miniata* genome and associated with specific droplets. Empty droplets were filtered using DIEM, ambient RNA was removed using SoupX, and doublets were predicted and removed using UMI counts. In total, 5857 droplets passed initial filtering and were used for subsequent clustering and analysis.

We clustered nuclei and identified marker genes for each cluster (Fig. 1A-B, Supp. Data S1). Clusters of poor quality nuclei retained through initial preprocessing were identified and removed, leaving 4372 nuclei among 25 clusters. Through inspection of marker gene sets, functional term enrichment analysis, and whole mount in situ hybridization, we were able to annotate these clusters (Fig. 1C-X, Supp. Data S1, S2), which include populations derived from all three developmental germ layers. Most cell clusters are in strong agreement with the cell types expected from previous morphological and molecular characterizations, and several clusters represent cell types previously unidentified in larval *P. min*. This data represents a comprehensive cell type atlas of normal *P. min* larva, resolving major ectodermal, endodermal, and mesodermal lineages alongside additional proliferative and specialized cell populations. Below is a brief summary of the populations identified. For more detailed information, please refer to the Supplementary Text.

**Figure 1.**
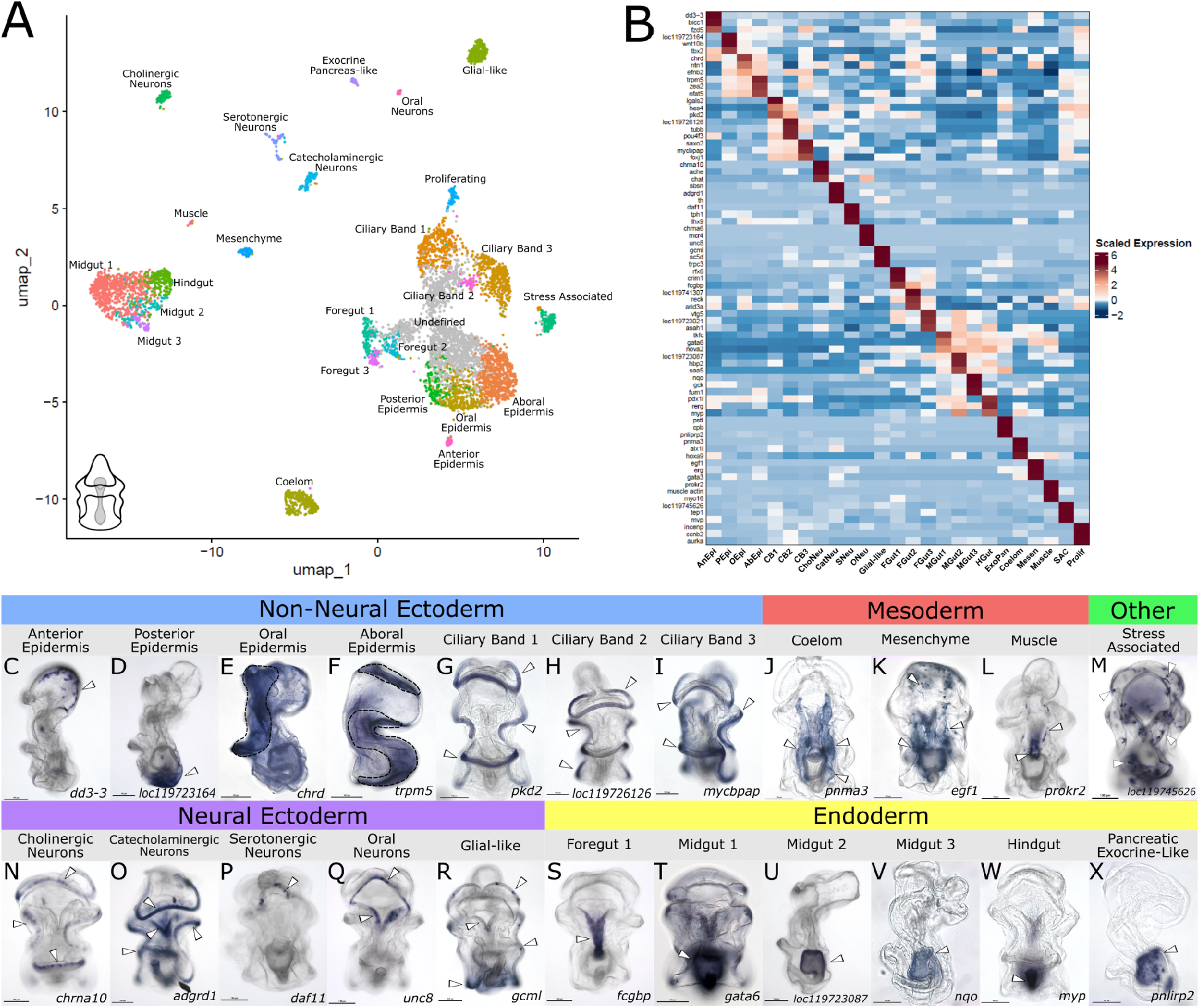
snRNA Atlas and Spatial Distribution of Intact Larval Cell Types. (A) UMAP projection of nuclei from intact 9 dpf *Patiria miniata* larvae. 25 distinct clusters are identified. Clusters are color coded and labelled. The undefined cluster not used in subsequent analyses is colored gray. (B) Heatmap of the most specific markers for each cluster. Marker genes are on the y-axis and clusters on the x-axis. Color represents the average expression of a gene within a cluster, scaled per row. AnEpi: anterior epidermis; PEpi: posterior epidermis; OEpi: oral epidermis; AbEpi: aboral epidermis; CB: ciliary band; ChoNeu: cholinergic neurons; CatNeu: catecholaminergic neurons; SNeu: serotonergic neurons; ONeu: oral neurons; FGut: foregut; MGut: midgut; HGut: hindgut; ExoPan: exocrine pancreas-like; Mesen: mesenchyme; SAC: stress associated cell; Prolif: proliferating. (C-X) In situ hybridizations for marker genes of 22 different clusters. White arrowheads indicate expression localization. Dashed lines for oral and aboral epidermis indicate expression localization.

Within the ectoderm, we identified distinct non-neural domains including multiple ciliary band subtypes and spatially patterned epidermal territories (Fig. 1C-I, Supp. Text), as well as diverse neural populations comprising cholinergic, catecholaminergic, serotonergic, oral, and glial-like cells with distinct molecular signatures and spatial organization (Fig. 1N-R, Supp. Text). Endodermal tissues were resolved into foregut, midgut, hindgut, and an exocrine pancreas-like population, with further subdivision revealing functional specialization related to digestion, absorption, and detoxification (Fig. 1S-X, Supp. Data S2, Supp. Text). Mesodermal clusters included coelomic, mesenchymal, and muscle populations, expressing transcription factors and immune-associated genes consistent with prior embryonic and larval studies (Fig. 1J-L, Supp. Data S1, Supp. Text).

Interestingly, we also detect the presence of proliferating cells based on the expression of genes associated with mitosis (Fig. 1A, 1B, Supp. Data S1, S2, Supp. Text). We identified the most specific markers for each cluster, yielding a set of highly specific markers for most clusters and supporting our clusters as biologically meaningful groupings (Fig. 1B, Supp. Data S3).

Notably, we also detect a previously unreported cell type enriched for genes associated with stress response (Fig. 1M, Supp. Data S2). These include immediate early genes such as the *transcription factor AP-1-like (jun) and Fos-related antigen 2-like (fosl2), as well as g*enes associated with MAPK signaling and the regulation of inflammation (Supp. Fig. S1A). Interestingly, markers indicating direct immune function are lacking in this cluster. We therefore annotate this cluster as stress-associated cells (SACs). SACs also express the eukaryotic major vault components, *telomerase associated protein 1* (*tep1*) and *major vault protein* (*mvp*), as well as the chromatin remodeler, *polycomb group protein Pc* (*pc*) (Fig. 1B, Supp. Fig. S1A). These cells are primarily located along both ciliary bands but can be detected in other parts of the ectoderm including the oral ectoderm (Fig. 1M), though we observe variability in the number and spatial distribution of these cells between different larval cultures, particularly along the AP axis, with cells frequently clustering in the anterior portion of the animal (Supp. Fig. S1B-G). These cells are sometimes observed displaying tear-drop morphologies or extending long projections, suggesting possible migratory behavior (Supp. Fig. S1Bi, S1Ci).

### Identification of normal and regenerating specific cell states

Using our intact larval data, we then investigated which intact larval cell states are gained, lost, or change broadly following bisection. To do this, we generated a snRNA-seq dataset from combined anterior and posterior fragments of *P. min* larvae collected 3 dpb–when wound proximal proliferation is first detected, and compared cluster composition with that of intact larvae. This dataset was preprocessed similarly to the intact larvae dataset, yielding 11143 filtered droplets in total. After clustering and classifying poor quality data as undefined, 25 distinct populations were observed, comprising 7098 nuclei (Supp. Fig. S2). Next, we integrated snRNA-seq data sets from intact and regenerating larvae using reciprocal principal component analysis (RPCA). To this end, we included the undefined clusters from both datasets to maximize the number of relationships available for integration. After integration, we performed clustering on the combined dataset, excluding undefined clusters to prevent inappropriate merging of predicted noise with genuine cell populations. This produced a final integrated dataset comprising 11470 nuclei and 27 integrated clusters (Fig. 2A).

**Figure 2.**
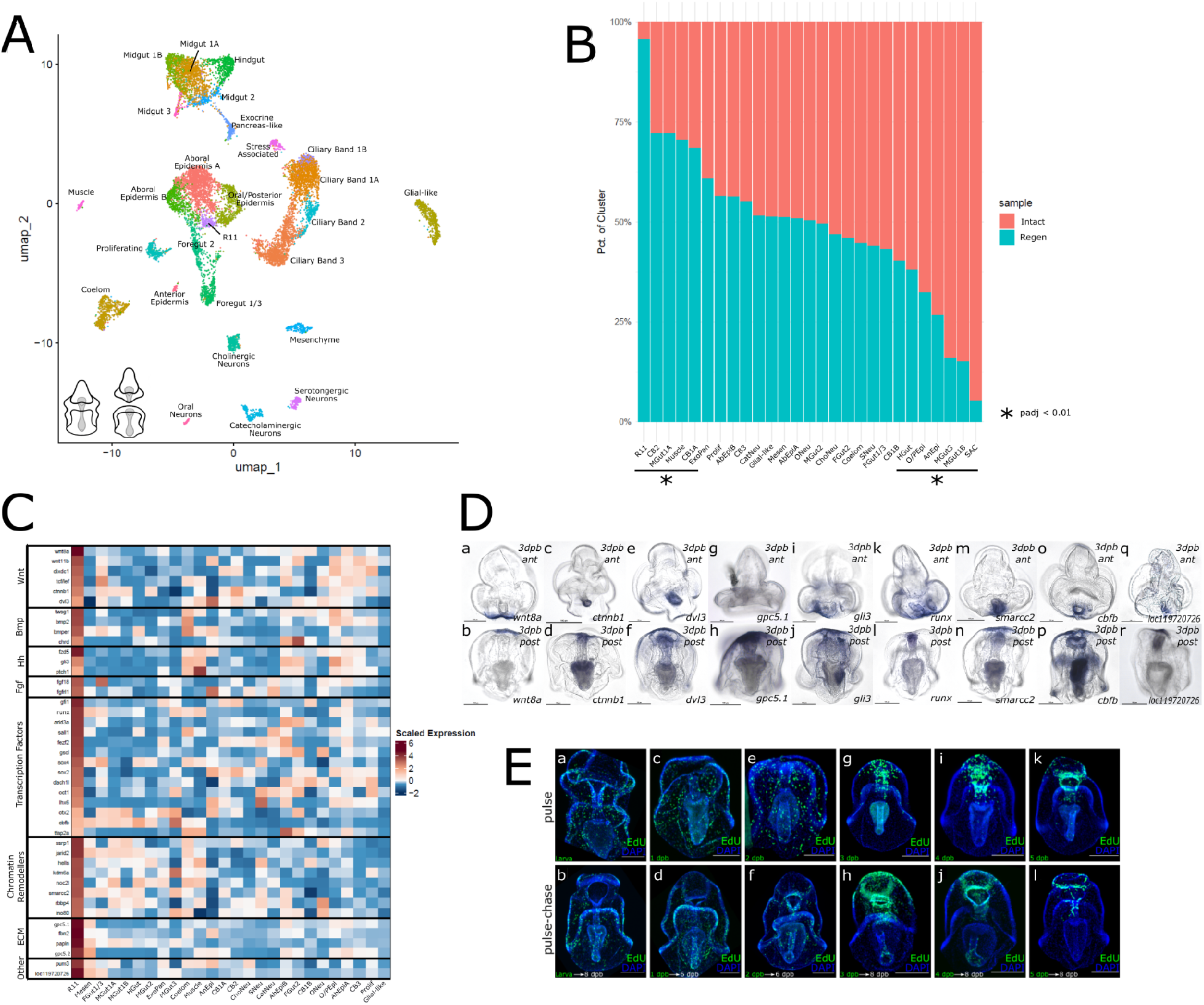
Integrated Cluster Sample Bias and Identification of the Blastema. (A) UMAP projection of nuclei integrated across intact 9 dpf larvae and 3 dpb regenerating fragments. 27 clusters are color coded and labeled according to the most similar 9 dpf dataset cluster identity, or 3 dpb identity in the case of R11. (B) Stacked bar plot depicting the relative abundance (y-axis) of intact larva (red) or 3 dpb larva (blue) nuclei within integrated clusters (x-axis), normalized for dataset size. Clusters are organized from mostly 3 dpb nuclei to mostly 9 dpf nuclei. Asterisks indicate significant enrichment for a condition (Fisher’s exact test, adjusted *p* < 0.01). (C) Heatmap of R11 marker genes grouped by functional similarly (y-axis) in different integrated clusters, excluding SACs (x-axis). Nuclei were filtered to only contain only those from 3 dpb larvae. Color represents average expression, scaled per row. Abbreviations for (B-C) similar to in Fig. 1. (Da-Dr) In situ hybridizations of R11 marker genes in 3 dpb anteriors (top row) and posteriors (bottom row), showing marker expression primarily at the wound site. (Ea-El) Larvae pulse-chased with EdU. Larvae pulsed with EdU (top row) show the localization of proliferating cells at specific time points. Chased EdU experiments (bottom row) show where cells proliferating at corresponding time points are localized at more advanced regenerative stages.

Measuring the joint membership of nuclei between integrated and unintegrated clusters, we observed that at least 50% of each unintegrated cluster maps to an integrated cluster, with most contributing over 90% (Supp. Data S4). This indicates that the unintegrated clusters are stable enough to maintain their identity after integration and enables us to assign annotations to integrated clusters based on intact larval cluster identities. One integrated cluster had little contribution from the intact larvae dataset (<10%, Fig. 2B), so we annotated this cluster using the regenerating larval cluster that contributes the most to it, R11.

We then investigated changes in cell type identity by determining whether any integrated clusters were enriched for intact or regenerating larval nuclei.

Most clusters show good mixing of nuclei from each sample, with roughly equal representation across most clusters (Fig. 2B). This shows that, in general, cell types do not change in relative abundance after bisection. Some clusters did, however, show significant enrichment for one sample (Fisher’s exact test, adjusted *p* < 0.01; Supp. Data S5). Those enriched for regenerating (i.e. 3 dpb) larval nuclei are R11, ciliary band 2, midgut 1A, muscle, and ciliary band 1A, and those enriched for intact larval nuclei are SACs, midgut 1B, midgut3, anterior ectoderm, oral/posterior ectoderm, and hindgut. Two integrated clusters, R11 and SACs, represent extreme cases, with 96% of R11 belonging to the regenerating larvae dataset and 95% of SACs belonging to the intact larvae dataset (Fig. 2B, Supp. Data S5).

### The blastema is a molecularly distinct population of cells induced by bisection

To investigate the regeneration-induced cluster, R11, we subset the integrated dataset to 3 dpb larval nuclei and identified marker genes. This cluster is enriched for GO terms associated with development and patterning and is marked by the expression of many components and regulators of pathways important for development including the WNT, BMP, Hedgehog, and FGF pathways (Fig. 2C, Supp. Fig. S3, Supp. Data S6, S7). We also detect expression of many transcription factors including *sox2*, *sox4*, *dach1l*, *runx*, and *core-binding factor subunit beta* (*cbfb*), which encodes a Runx transcriptional co-regulator (Chuang et al., 2013), as well as several chromatin remodelers.

Furthermore, we detect strong expression of the ECM regulator, *glypican-5-like* (*gpc5*), which can also regulate WNT signaling (Yuan et al., 2016).

In situ hybridization reveals that all markers tested for R11 are expressed at the wound site in 3 dpb larvae for both anterior and posterior fragments with the exception of *gpc5* (Fig. 2Dg). This domain coincides with the site of rapid proliferative expansion observed around 3 dpb, which colocalizes with the R11 markers *runx*, *dach1l*, *gli3*, *mcm2*, *sox2*, and *sox4* (Cary et al., 2019; M. Zheng et al., 2022). This site is predicted to be functionally equivalent to the blastema, which is seen in several regenerative systems and that we will define as a mass of proliferating cells contributing directly to the tissues of the regenerate.

To test the functionality of the predicted blastema in *P. min* regeneration, we performed a pulse-chase experiment using EdU to determine the fate of proliferating cells during regeneration in larvae. In posterior fragments, cells labelled with EdU prior to 3 dpb are distributed throughout the body and do not substantially contribute to the regeneration of anterior tissues. Cells labeled at 3 dpb are concentrated at the proximal wound site and importantly, through pulse chase we see that these later contribute to the regenerated tissues (Fig. 2E).

Notably, cells labeled after 3 dpb contribute progressively less to the regenerated anterior, indicating that cells destined to form the regenerated tissues are committed primarily in this critical time window (Fig. 2Ei-El). This confirms the regenerative blastema’s functionality in P. min and supports the annotation of R11 as such.

Interestingly, blastema marker expression is observed in non-overlapping domains within the wound site, confirming previous reports (Cary et al., 2019). Expression is detected in the epidermis, as with *wnt8a*, the foregut, as with *runx*, and the regenerating bilateral coelom, as with *gli3* (Fig. 2Db, Dj, Dl). This suggests a level of heterogeneity not captured at the clustering resolution used for the integrated dataset, although the unified clustering of these cells indicates a broader pattern of shared gene expression within this population.

### Blastema Formation is Controlled by MAPK Signaling and is Associated with the Disappearance of SACs

We then investigated the functional significance of cell signaling on the formation of the blastema. Specifically, we perturbed the MAPK/ERK signaling pathway using the pharmacological inhibitor, U0126, which has been shown to impair blastema formation in several systems (Tasaki et al., 2011; Suzuki et al., 2007). We found that treatment with 15 μM for 24 hours is sufficient to prevent ERK phosphorylation in 1 hour post-bisection (hpb) larval fragments without noticeable defects in wound healing (Fig. 3A-E) and also prevents wound localized proliferation in 3 dpb fragments, indicating impaired blastema formation (Fig. 3F-G).

**Figure 3.**
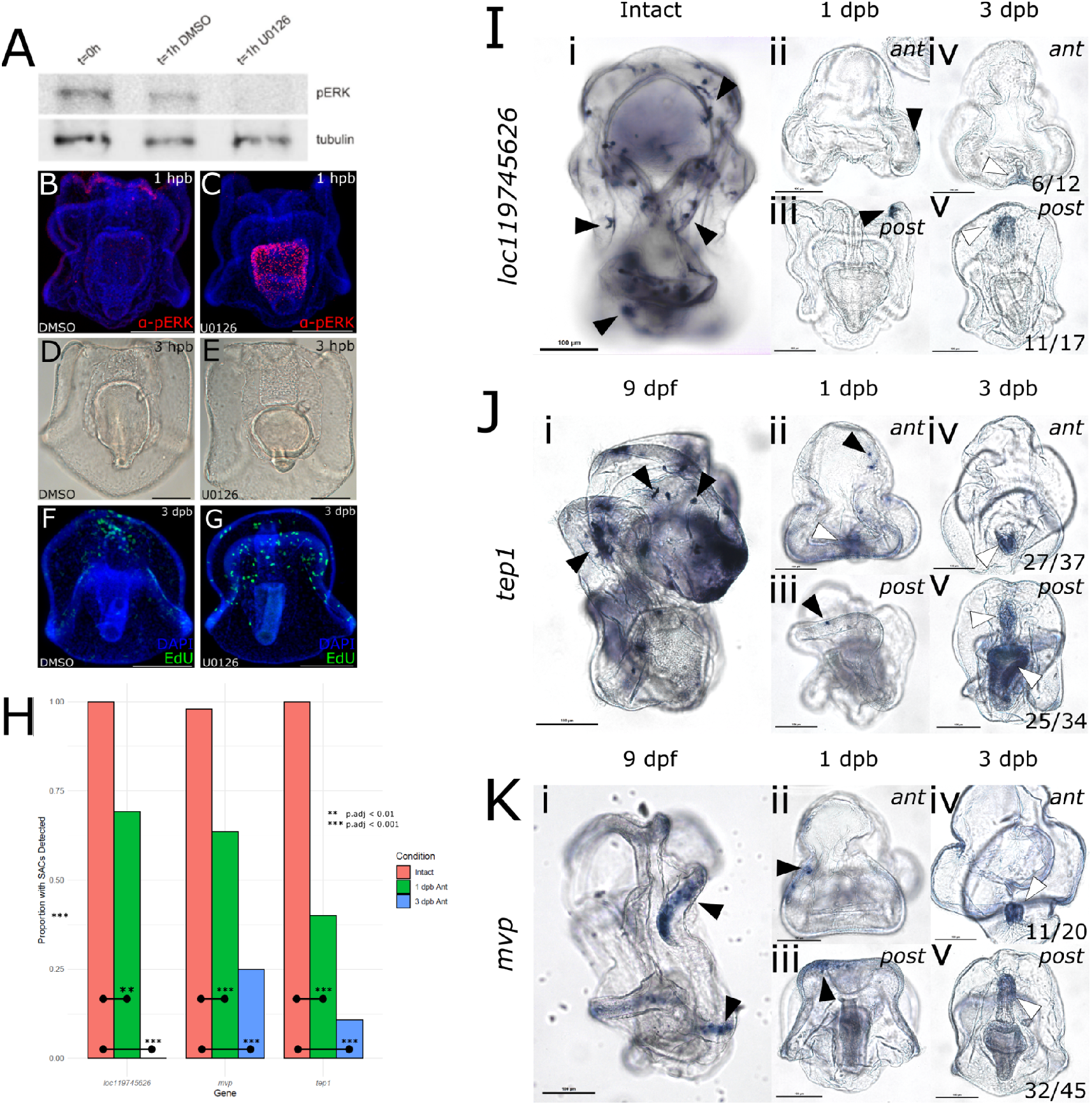
SAC Marker Expression Shifts From Epidermal SACs to the Pharyngeal Blastema. (A) Western blot of pERK and tubulin from embryos treated with either DMSO or U0126. (B-C) pERK localization in control or U0126 treated posterior 1 dpb fragments, showing loss of wound-localized pERK in treated samples. (D-E) Bright field images indicate no effect on wound closure in control or U0126 treated 3 hpb posterior fragments. (F-G) EdU labeling of 3 dpb posterior fragments treated with DMSO or U0126. (I-K) In situ hybridizations of SAC markers in intact larvae, 1 dpb larval fragments, and 3 dpb larval fragments. Black arrowheads indicate signal in predicted SACs. White arrowheads indicate signal in the regenerating endoderm, with values in 3 dpb panels indicating the proportion of fragments with signal detected in the regenerating endoderm. (H) Quantification of SAC loss after bisection. For all markers, we observe significant decrease in the number of anterior fragments with SACs after bisection (adjusted *p* < 0.01 pairwise chi-squared test for equality of proportions)

We also followed up on the apparent loss of SACs that was observed at 3 dpb in our snRNA-seq data (Fig. 2B). All SAC markers that we tested for exhibited a sharp and significant decrease in punctate epidermal expression as early as 1 dpb, continuing into 3 dpb (Fig. 3H-K, adjusted *p* < 0.01 pairwise chi-squared test for equality of proportions). Interestingly, this is concurrent with an increase in expression in the regenerating endoderm of over half of all samples (Fig. 3I-K). This is the same time and place where blastema markers are expressed (Fig. 2D), suggesting that SAC marker expression has shifted to the blastema.

### Identification of Regeneration-Responsive Elements

Having identified the blastema as a regeneration-specific cell type contributing to regeneration, as well as the transcriptional state defining this cell type, we then sought to understand how this state is regulated at the cis-regulatory level through enhancers. As many genes identified in the blastema are related to developmental patterning, we were particularly interested in whether these genes are regulated by regeneration-specific regulatory mechanisms, such as wound-induced signaling, or are co-opted from embryogenesis. To do this, we generated ATAC-seq data sets for intact larvae and combined anterior and posterior fragments from 3 dpb larvae to identify regeneration-responsive elements (RREs), which we define as those with differential accessibility between conditions. We also processed embryos collected at 1-, 2-, and 3 dpf, representing the blastula, mid-gastrula, and late-gastrula stages, respectively, to determine whether any RREs are shared with embryonic stages. In addition, we included an early wounding time point at 1 hpb to examine how early RREs change accessibility after bisection.

Each time point was represented by 2 biological replicates, each with 2 to 3 technical replicates. We used read insert length, transcription start site enrichment (TSSE), and replicate concordance as measures of data quality (Sup Fig. S4). Read length displays weak but detectable periodicity corresponding to nucleosome length, which is acceptable, and reads show high TSSE, which is ideal (Supp. Fig. S4A-B). Replicate concordance was assessed using principal component analysis (PCA), which shows high similarity between technical replicates (Supp. Fig. S4C). Biological replicates are notably different from each other, but both replicates take the same trajectory along the PCA plot over time, indicating replicable changes in chromatin dynamics over the time series. Taken together, these support the quality of our dataset.

Consensus open chromatin regions (OCRs) were identified by filtering for regions present in at least two technical replicates, which yields 106756 OCRs. Accessibility levels for these OCRs were then compared between time points.

62.1% show significant accessibility change between at least two time points (adjusted *p* < 0.01 Wald test, fold change > 1.5), revealing a highly dynamic chromatin landscape. Between intact and 3 dpb regenerating larvae, 1981 differentially accessible regions were detected, which we denote as RREs (Fig. 4A. Supp. Data S9). 98 total OCRs are differentially accessible between intact and 1 hpb larvae, 9 of which increase in accessibility after bisection, which we predict as wound-responsive enhancers (Supp. Fig. S5B). Notably, only 13 RREs are also wound-responsive. However, these have very weak signal in intact larvae and almost no signal at any other time point (Supp. Fig. S5A). Therefore, these are intact larva-specific and are not necessarily RREs activated early.

**Figure 4.**
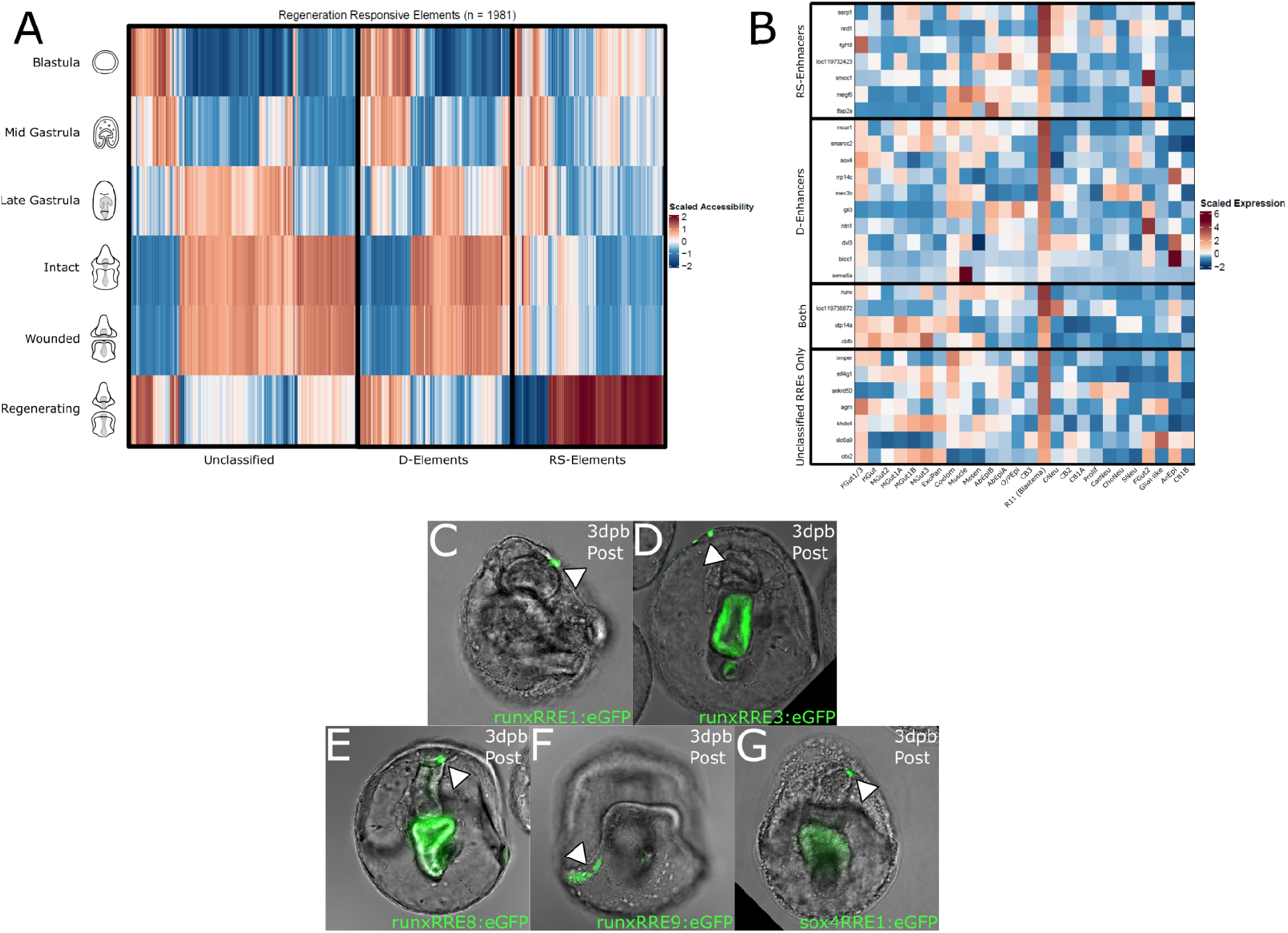
Regeneration-Responsive Elements Function in the Blastema. (A) Heatmap of OCRs (x-axis) with significantly different accessibility (Wald test, adjusted *p* < 0.01) between intact and 3 dpb regenerating larvae. Samples on the y-axis and ordered temporally. OCRs are clustered using Pearson correlation. Regeneration-specific (RS-elements), developmental (D-elements), and unclassified RREs are enclosed in black borders. Color represents accessibility, scaled per column. (B) Heatmap of the expression of different blastema markers (y-axis) predicted to be under the regulatory control of different RRE subsets and grouped accordingly. Markers are predicted to be regulated by RS-elements, D-elements, both, or neither. All of these may also include regulation by unclassified RREs. Different integrated clusters are shown on the x-axis, and color represents average marker expression in that cluster, scaled per row. Data was filtered to only contain 3 dpb larvae nuclei. 4 genes are predicted to be regulated by regeneration-specific and developmental RREs. Cluster name abbreviation conventions are similar to in Fig. 1. (C-G) Fluorescent microscopy images showing the expression of an eGFP reporter under the regulatory control of different RREs. All constructs drive eGFP expression (white arrowhead), and 4 drive expression at the wound site. Autofluorescence is observed in the midgut in the images for *runxRRE3:eGFP, runxRRE8:eGFP, and sox4RRE1:eGFP* injected animals.

We predicted downstream functionality of the 9 wound-responsive enhancers by associating them with nearby genes, which include *early growth response protein 1-B* (*egr1b*) and *E74 like ETS transcription factor 2* (*elf2*), which could have pioneer transcription factor activity (Gehrke et al., 2019; Xiao et al., 2024), and a chromatin remodeler, *histone deacetylase 11-like* (*hdac11*) (Supp. Data S10).

RREs display a wide range of temporal dynamics across the time series (Fig. 4A). For instance, some RREs exhibit extreme accessibility at 3 dpb (fold change > 1.5 relative to all other time points), which we classify as regeneration-specific RREs (RS-elements). Other RREs return to accessibility levels observed during embryogenesis (fold change < 1.2 between 3 dpb and any embryonic time point), which we classify as developmental RREs (D-elements). In total, we detected 564 RS-elements, 566 D-elements, and 850 unclassified RREs. Of the 566 D-elements, 192 are blastula-like, 265 mid-gastrula-like, and 208 late-gastrula-like RREs, with only 90 of these (15.9%) shared with more than one embryonic time point (Supp. Fig. S7). These data indicate that there is no enrichment for the use of regeneration-specific or developmental regulatory mechanisms at 3 dpb, and that globally there is no evidence that regeneration activates D-elements in a temporal order mirroring development.

### Regulation of Blastema Gene Expression Patterns

We then associated opening RREs (i.e. enhancer RREs) with nearby genes and filtered these enhancers to those associated with significant blastema marker genes (adjusted *p* < 0.05 Wald test, Supp. Data S7, S9). In other words, we searched for enhancers that are predicted to increase activity at 3 dpb and are near genes associated with the blastema. In total, we found 47 enhancer RREs associated with 28 blastema markers (Fig. 4B). Most of these marker genes are predicted to be regulated by either RS-enhancers or D-enhancers, and many fall into important functional groups, such as transcription factors, chromatin modulators, and signalling pathway components and regulators (Fig. 2C). RS-enhancers are associated with *ssrp1*, *fgf18*, and *tfap2a*. D-enhancers are associated with *smarcc2*, *sox4*, *gli3*, and *dvl3*. *Otx2* is associated with enhancer RREs classified as neither RS- nor D-enhancers, and 4 markers, including *runx* and *cbfb*, are predicted to be regulated by both RS- and D-enhancers. Interestingly, of 712 total genes regulated by enhancer RREs, only 31 (4%) are co-regulated by both RS- and D-enhancers (Supp. Fig. S8, Supp. Data S9), suggesting that only a relatively small proportion of genes, including *runx* and *cbfb*, are able to integrate the regulatory activity of these different enhancer sets and potentially coordinate their downstream effects.

We then validated a subset of enhancer RREs predicted to regulate blastema expression by cloning them into reporter constructs containing an H2B minimal promoter upstream of eGFP. We then tested for their function in regenerating larvae. Constructs were injected at the 1-cell stage, and larvae negative for eGFP were bisected and allowed to regenerate for 3 days, reducing the chance of false-positive signals carried over from before bisection. Using this method, we confirmed the function of 4 enhancer RREs predicted to regulate *runx* and a single enhancer RRE predicted to regulate *sox4* (Fig. 4C-G). This set includes both RS-enhancers, including *runxRRE1, runxRRE3,* and *runxRRE9*; and D-enhancers, including *runxRRE8* and *sox4RRE1*. *Runx*RRE1 and *runx*RRE3 both drive reporter expression in what appears to be the wound epidermis in posterior fragments (Fig. 4C-D). *Runx*RRE8 and *sox4*RRE1 activity is found in the regenerating mouth and foregut, respectively (Fig. 4E, G).

*Runx*RRE9 activity is found in the posterior ciliary band away from the wound site (Fig. 4F). These data confirm enhancer regulatory activity in the blastema as well as tissues distal to the wound site, and support our ability to identify CREs that drive gene expression during regeneration in *P. min* larvae.

We next predicted the transcription factors (TFs) regulating gene expression through the 4 enhancer RREs confirmed to function in the blastema. To do this, we performed TF footprinting analysis using TOBIAS, which uses accessibility information from ATAC-seq to indirectly measure protein occupancy within DNA motifs and to calculate differences in average occupancy between conditions (Bentsen et al., 2020). This analysis requires known binding motifs for each TF, and for this we used the vertebrate profiles from JASPAR. The full results for this analysis, including all increases and decreases in TF binding within RREs, can be found in Supplementary Data S11. We then filtered these to include only binding increases between 3 dpb and intact larvae in TF motifs either marking the blastema or known to be activated by pERK, revealing extensive binding changes within RREs predicted to control blastema gene expression (Supp. Data S12).

Within the 4 functionally validated blastema enhancers, we found increased binding of Tcf7, Sox14, Runx, and Elk within runxRRE3, of Fos, Jun, Srf, Tfap2a, and Gfi1 within runxRRE8, and of Creb, Fos, Jun, Myc, and Runx within sox4RRE1 (Supp. Data S12). These results indicate that Runx is able to autoregulate its own expression through regeneration-specific enhancer elements. Furthermore, Runx can potentially drive *sox4* expression by interacting with developmental enhancers. We also note that both D-elements (runxRRE8 and sox4RRE1) have more sites with increased binding of various pERK targets, whereas the runxRRE3, an RS-element, only has one Elk site with increased binding, hinting at separate regulatory activity of RS- and D-elements in terms of TF binding, similar to how these elements regulate largely separate groups of genes (Supp. Fig. S8).

To assess the generalizability of these findings, we ran footprinting analysis on all RS- and D-elements and compared the average binding change for each motif between these sets. Average binding change shows a positive but weak correlation between RS- and D-elements (R^2^ = 0.321, Supp. Fig. S9D), indicating differential transcription factor usage. In fact, looking at the motifs with significant change in average binding score (−log_10_(adjusted *p*) > 95th percentile, absolute differential binding score > 95th percentile), only 3 motifs show either a significant increase or decrease within both RS- and D-elements, those being Runx1, Runx2, and Bcl11b (Supp. Fig. S9A-C, Supp. Data S13). Because the JASPAR binding motif for Bcl11B, which is known to form a complex with Runx (Sidwell & Rothenberg, 2021), is nearly identical to that of Runx, and because there is only 1 *runx* homolog in the *P. min* genome, we effectively interpret these results as Runx being the only factor with a significant binding increase in both RS- and D-elements. Furthermore, of the 47 enhancer RREs associated with blastema genes, 27 have Runx1, Runx2, or Bcl11b motifs with increases in binding during regeneration, which are associated with 21 of 28 enhancer RRE-associated blastema markers (differential binding score > 1, Supp. Data S11). These results, coupled with the fact that *runx* is associated with both RS- and D-enhancers, support a general role for Runx as an integrating component of regeneration-specific and developmentally shared regulatory mechanisms, notably in the blastema, allowing one mechanism to feed regulatory input into *runx* which then can directly influence the other mechanism through binding of Runx protein.

## Discussion

In this study, we sought to identify the cells that respond to wound-induced signals following bisection, define their molecular identities, and reconstruct the gene regulatory networks (GRNs) that connect wounding to the reactivation of developmental and/or regeneration-specific programs. We further asked how regeneration-specific and developmental GRNs interact to re-establish the cell lineage trajectories required to replace lost tissues. A summary of our findings and predicted model is presented as a GRN model in Figure 5 and Supplementary Data S13.

**Figure 5.**
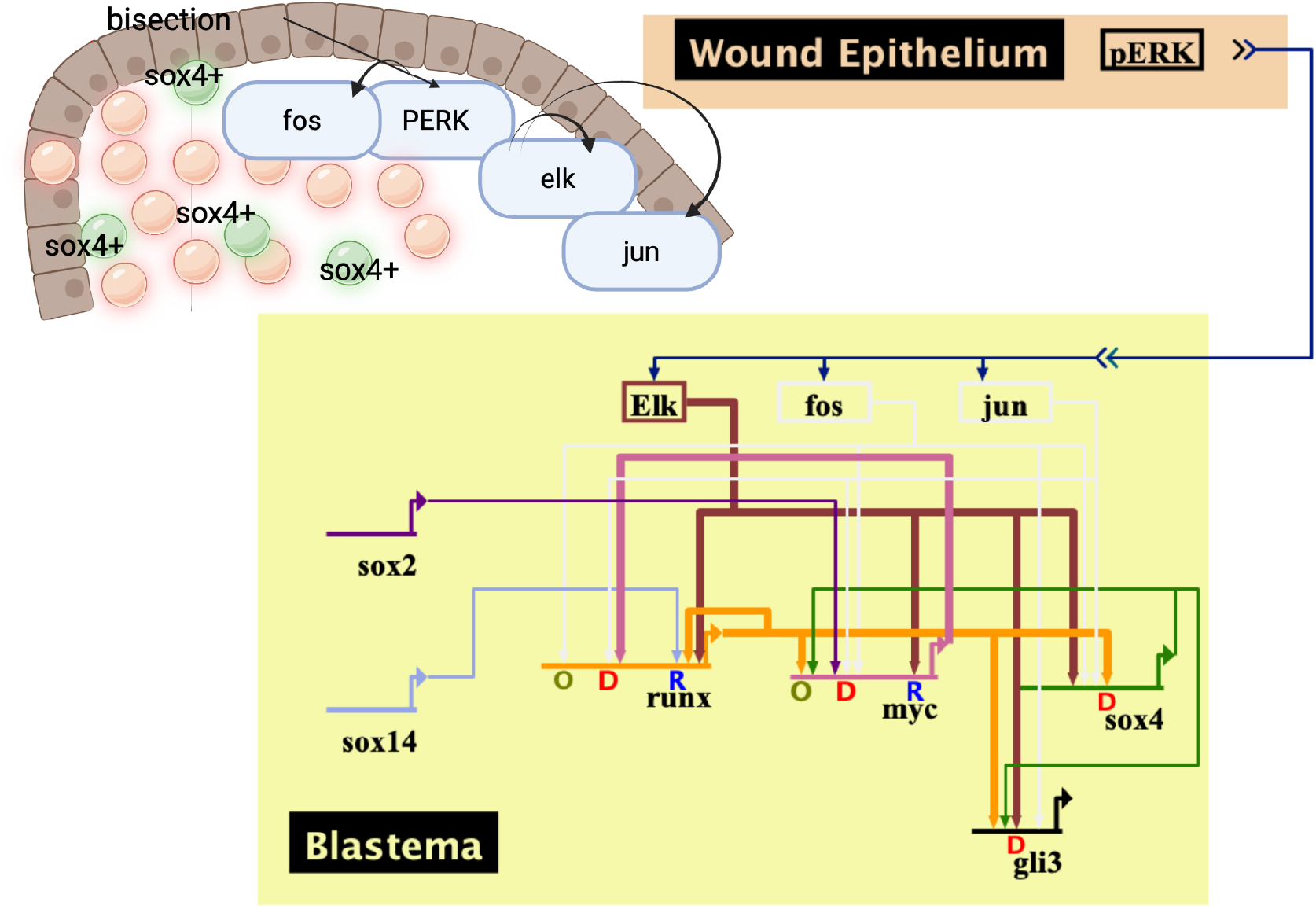
Model GRN driving expression of developmental genes during blastema formation. Following bisection, ERK protein is phosphorylated in the wound epithelium, without which blastema formation is inhibited. This is predicted to activate Elk, Fos, and Jun protein in blastema cells as depicted by connections with double arrowheads. This increases binding of ELK, Fos, and Jun to enhancer elements controlling gene expression in the blastema. Direct regulatory interactions are shown as arrows between genes and enhancers, which are labelled as regeneration-specific (R), developmental (D), or other (O). This ultimately leads to the activation of a GRN controlling the expression of multiple developmental transcription factors, including *sox4*.

By integrating snRNA-seq datasets from intact larvae and larvae at 3 dpb, we examined how cell states change following injury, including whether specific states are lost or gained and whether transcriptional changes occur globally or are restricted to defined populations. This analysis revealed a regeneration-induced molecular identity corresponding to the blastema in larval sea star regeneration. Importantly, most cell states are equally represented before and after regeneration, indicating that cell identity change in response to wounding is restricted rather than global. Markers of this blastema population further suggest that it functions as a signaling center, potentially coordinating multiple developmental pathways, consistent with an organizer-like role during regeneration.

A central insight from our integrated transcriptomic and chromatin accessibility analyses is that regeneration in *P. min* is driven by distinct classes of regeneration-responsive enhancers that link wounding signals to GRN activation. These enhancers include both regeneration-specific elements and developmentally reused elements, whose activities are coordinated through the transcription factor Runx. Together, this regulatory architecture provides a mechanistic framework for integrating regeneration-specific and developmental programs during tissue reconstruction.

Our single-nucleus atlas of intact larval *P. min* reveals several previously undescribed cell populations. Among these, the identification of stress-associated cells (SACs) was unexpected and raises several questions. No analogous cell type has yet been described in other echinoderms, and the functional role of these cells remains unclear. Based on their expression of *ap-1* and MAPK signaling components, SACs may be poised to respond to cytokines or other environmental stress signals and relay this information to surrounding tissues. The expression of *mvp* and *tep1*, components of the major vault complex, is also notable. Major vaults are conserved across eukaryotes but remain poorly understood; in echinoderms, they have only been described in the context of nuclear and nuclear envelope localization (Hamill & Suprenant, 1997).

SACs were notably undetectable in our regenerating larval snRNA-seq data sets. Given that TUNEL staining does not reveal a significant increase in apoptosis until the onset of blastema proliferation (Cary et al., 2019), well after the reduction in SACs at 1 dpb, it is unlikely that these cells are eliminated by apoptosis. Instead, SACs may persist following bisection but lose their stress-associated transcriptional identity, and could either contribute directly to the blastema or provide signals that facilitate its formation, although these possibilities remain to be tested.

Many genes identified as markers of the regeneration-induced blastema state are expressed within the same broad population of cells at the wound site. However, although our snRNA-seq analysis classifies the blastema as a single cluster, in situ hybridization reveals non-overlapping expression patterns among individual markers, suggesting cellular heterogeneity that is not fully resolved at the depth of our dataset. Notably, the same sets of blastema markers are expressed at both anterior- and posterior-facing wound sites, indicating that these genes are not directly involved in axial patterning. For example, we observe wnt8a expression at the anterior end of posterior fragments, despite its established role in posterior fate specification during embryogenesis (McCauley et al., 2013; Hikasa & Sokol, 2013).

We detect expression of components of several major signaling pathways, including Wnt and Hedgehog, suggesting that the *P. min* blastema may function as an organizer that can autonomously direct patterning during regeneration.

This role is reminiscent of the wound epidermis in vertebrate blastemas, which similarly acts as a signaling hub during regeneration (C Aztekin et al., 2019; Stoick-Cooper, Moon, et al., 2007; E. M. Tanaka, 2016). The classification of epidermally derived blastema-like structures, as observed here in larval *P. min* and in adult echinoderms , has been contentious, in part because traditional definitions of blastemas emphasize mesenchymal sources of proliferating cells (Czarkwiani et al., 2016; Moss et al., 1998; Ben Khadra et al., 2018). Our findings suggest that a shared feature of blastemas may instead lie in their signaling capacity rather than in the cellular origin of proliferative cells. Emphasizing signaling over mesenchymal contribution may help explain variability in blastema characteristics across single-cell studies of regeneration (Gerber et al., 2018; Leigh et al., 2018; Storer et al., 2020; Wang et al., 2020). Under this framework, the blastema in larval *P. min*, as well as epithelial blastemas in adult sea stars and brittle stars, may arise through the concentration of proliferative and signaling functions within epithelial populations.

To define the regulatory basis of the enhancer-driven framework described above, we generated ATAC-seq time-series datasets spanning early development (blastula, mid and late gastrula), larval stages, and regeneration. We identified thousands of potential cis-regulatory elements that change in accessibility during regeneration, which we term regeneration-responsive elements (RREs). By associating RREs with nearby genes and referencing these to genes expressed in the blastema, we identified multiple candidate enhancers likely to regulate blastema-specific transcription. These RREs can be classified into regeneration-specific (RS) and developmental (D) elements, which are largely associated with distinct gene sets and differ in their transcription factor binding profiles. It is important to note, however, that this analysis primarily focuses on elements that increase in accessibility and may therefore miss repressive elements that close during regeneration.

These analyses identify Runx as a central regulatory node linking regeneration-specific and developmental enhancer activity. Runx binding sites are enriched among both RS- and D-elements, and most show increased accessibility during regeneration. Enrichment of Runx motifs has also been observed in other regenerating systems, suggesting a conserved role for Runx in regeneration (Gehrke et al., 2019; Goldman et al., 2017; Hughes & Woollard, 2017; Kawaguchi et al., 2024). Many transcription factors are associated with Runx-interacting RREs, indicating that Runx may function as an important regulator coordinating regeneration-associated GRNs. This includes *runx* itself, consistent with autoregulatory behavior previously described in other regenerative contexts (Goldman et al., 2017).

Within the blastema, we provide evidence that *sox4* is regulated by Runx through developmentally reused enhancers. This finding is particularly significant given our previous demonstration that *sox4*⁺ cells arise *de novo* during regeneration.

Specifically, *sox4* displays regeneration-specific expression patterns, including expression in non-neural cell types and in cells lacking *sox2*, in contrast to its canonical role during embryonic neurogenesis (M. Zheng et al., 2022). Here, we identify a gene regulatory network that accounts for the emergence of this regeneration-specific *sox4^+^* expression state, linking injury-induced enhancer activation to *sox4* expression via Runx-dependent regulation. Together, these results indicate that *sox4* is activated during regeneration through enhancers shared with development but deployed in novel regulatory contexts, providing a mechanistic explanation for the *de novo* appearance of *sox4*⁺ cells.

Ultimately, our results support a working model of how regeneration activates developmental gene-expression programs. Bisection of *P. min* larvae leads to the specification of a blastema population that shares a common transcriptional profile while retaining internal heterogeneity. Stress-associated cell identities are suppressed following injury and may contribute indirectly to blastema formation, although this remains unresolved. Regeneration-responsive enhancers drive gene expression in the blastema through both regeneration-specific and developmentally reused mechanisms. Both enhancer classes converge on Runx, which coordinates downstream regulatory programs, including putative activation of developmental genes such as *sox4*, providing a regulatory explanation for the emergence of regeneration-specific cell states.

Together, these findings address how wound-induced signals specify regenerative cell states and reveal how regeneration-specific and developmentally reused gene regulatory networks are coordinated to rebuild lost tissues. By linking injury-induced enhancer activation to lineage re-establishment through Runx-dependent regulatory logic, this work provides a mechanistic framework for understanding how regeneration integrates novel and developmental programs.

## Methods

### Larval Cultures

#### Fertilization and Care

Larval cultures were fertilized as previously described (Cheatle Jarvela & Hinman, 2014). Briefly, gonads were collected from adult *Patiria miniata* females and shredded to release oocytes. These were then incubated in 1-methyladenine diluted in artificial sea water (ASW) at a final concentration of 10 μM for 45-90 minutes at 15 °C to mature. After maturation, gonad tissue and small unviable oocytes were filtered out using 200 μm and 100 μm mesh screens, respectively, and sperm from adult males was added to the oocytes. After fertilization, which takes 5-10 minutes, sperm were filtered using 100 μm mesh screens. Fertilized embryos were maintained in ASW at 15 °C and allowed to develop until needed. Water was changed on day 3 and every other day after until animals were needed for experiments. On day 4, early larvae were fed algae (*Rhodomonas lens*).

#### Microinjection

Microinjections were performed as previously described (Cheatle Jarvela & Hinman, 2014). After fertilization, embryos used for microinjection were incubated in ASW pH 4.0-4.2 for 5 minutes. For 2 of these, embryos are passed back and forth through a 200 μm mesh screen. Acid treated embryos are then rowed on the lids of new 60 x 15 mm polystyrene Petri dishes and injected using thin walled borosilicate glass capillaries with filament (Outer diameter 1.0mm, Inner Diameter 0..78mm, Overall length 10cm; Sutter Instruments) pulled using a Model P-97 Needle Puller (Sutter Instruments). Injection solutions were for DNA reporter constructs: 120mM KCl, 2x Rhodamine dextran, 5 ng/μL reporter construct DNA, 5 ng/μL HindIII digested genomic DNA.

#### Bisection

Animals were bisected at bipinnaria larval stage between 7 and 9 days-post-fertilization (dpf). Larvae were bisected midway between the anteroposterior axis through the foregut and bilateral coeloms. Bisected larvae were kept at 15 °C in 60 x 15 mm polystyrene Petri dishes at a concentration of around 75 larvae, or 150 larval fragments, per dish. Bisected animals were given fresh changes of ASW every day until they were processed for experiments.

#### snRNA-seq

Single nucleus RNA sequencing (snRNA-seq) experiments were performed using 9 dpf intact larvae and 3 dpb combined regenerating anterior and posterior larval fragments.

#### Nuclear Isolation

Nuclear isolation was performed as previously described with modifications (Meyer et al., 2023). Briefly, animals were hand-counted and resuspended in cold 10 mL filter sterilized ASW. Samples were centrifuged at 220 *g* and 4 °C for 5 minutes. ASW was discarded, and samples were resuspended in 10 mL HB buffer (15 mM Tris pH 7.4, 10% filter sterilized sucrose, 15 mM NaCl, 60 mM KCl, 0.2 mM EDTA, 0.2 mM EGTA, 2% BSA, 0.02 U/µL SUPERase-in RNAse inhibitor [Invitrogen], and cOmplete protease inhibitor [Roche] at a concentration of 1 tablet per 25 mL) and dounce homogenized on ice. Samples were then transferred to a fresh 15 mL conical tube and centrifuged at 3500 *g* and 4 °C for 5 minutes, after which the supernatant was discarded. Samples were resuspended in 1 mL of 1x PBS with 0.1% Triton X-100, 2% BSA, and 0.02 U/µL SUPERase-in RNAse inhibitor and then filtered through a 40 μM Flowmi Cell Strainer (Scienceware). This was then centrifuged at 3500 *g* and 4 °C for 5 minutes, the supernatant was removed, and nuclei were resuspended in 1 mL of 1x PBS with 0.1% Triton X-100, 2% BSA, and 0.02 U/µL SUPERase-in RNAse inhibitor. Nuclei were then counted by staining an aliquot with Trypan Blue and loading this onto a Countess II Automated Cell Counter (Thermo Fisher).

#### Library Prep and Sequencing

Libraries were prepared using the Chromium Next GEM Single Cell 3’ Reagent Kit v3.1 RevD (10x Genomics) according to the manufacturer’s protocol. Nuclei were loaded at a concentration of 1000 nuclei/µL onto a Chromium X Controller (10x Genomics) to be partitioned for a targeted recovery of 5000 nuclei. Partitioned nuclei-containing droplets were then incubated in a reverse transcription reaction. cDNA was then amplified and size selected using SPRIselect beads (Beckman Coulter). cDNA libraries were then fragmented, end repaired, A-tailed, ligated to adapters, and provided with sample indices.

Concentration of final libraries was measured using a Qubit Fluorometer (Invitrogen) and sequenced on NovaSeq 6000 high-throughput sequencer using a PE150 configuration for a target yield of 83 GB of data per sample, which equates to roughly 277 million reads per sample or 56 thousand reads per nucleus.

#### Data Analysis

Read data returned from sequencing was mapped using Cell Ranger 9.0.0 (G. X. Y. Zheng et al., 2017) to a reference protein coding transcriptome generated using the Pmin_3.0 *Patiria miniata* genome (GCF_015706575.1). Data matrices were imported into the R package, Seurat V5 (Hao et al., 2024), followed by empty droplets filtering using DIEM (Alvarez et al., 2020), ambient RNA contamination removal using SoupX (Young & Behjati, 2020), and doublets were removal using UMI cutoffs based on a prediction of 0.8% doublet rate per 1000 droplets.

Through Seurat, preprocessed data was normalized using SCT normalization, highly variable features were selected and used for dimensionality reduction using PCA, K-nearest neighbor graphs were constructed, and droplets were clustered using the Louvain algorithm. Visualization of clusters was performed using UMAP dimensionality reduction. Clustering was performed iteratively until an appropriate clustering structure and resolution were achieved. Cluster marker genes were identified using the Wilcoxon rank sum test and used to inform cluster annotations.

Individual datasets were integrated using reciprocal PCA through Seurat.

Datasets were individually normalized, shared highly variable features were used for PCA, and integration anchors were identified. Datasets were integrated using these anchors, and the resultant integrated dataset was used for clustering, as with the unintegrated datasets.

#### ATAC-seq

ATAC-seq was performed using a modified version of the Kaestner lab’s 2019 OMNI-ATAC-seq protocol. Samples were collected from 26 hours-post-fertilization (hpf) blastula, 42 hpf mid-gastrula, 72 hpf late gastrula, 7 dpf intact bipinnaria larva, 1 hour-post-bisection (hpb) combined anterior and posterior larval fragments, and 3 dpb combined anterior and posterior larval fragments. Each sample contains two biological replicates, where one culture was used for all time points in a biological replicate, and two to three technical replicates each, which were generated at the transposition reaction step.

#### Nuclei Isolation

Animals were hand counted and resuspended in 10 mL cold filter sterilized ASW. Animals were then centrifuged at 220 *g* and 4 °C for 2 minutes. ASW was removed, fresh cold ASW was added for a second wash, animals were centrifuged again. ASW was removed, and animals were incubated in 500

Animals were incubated in 500 µL lysis buffer (10 mM Tris-HCl pH 7.5, 10 mM NaCl, 3 mM MgCl_2_, 0.1% Tween-20, 0.1% NP-40, 0.01% Digitonin) on ice for 3-5 minutes. Samples were homogenized via pipetting, filtered through a 40 μM Flowmi Cell Strainer (Scienceware), and resuspended in 5 mL wash buffer (10 mM Tris-HCl pH 7.5, 10 mM NaCl, 3 mM MgCl_2_, 0.1% Tween-20). Samples were centrifuged at 500 *g* and 4 °C for 10 minutes, supernatant was removed, samples were resuspended in 1 mL cold PBTB (1x PBS, 0.1% Tween-20, 2% BSA) and filtered once more through a 40 μM Flowmi Cell Strainer. An aliquot of nuclei were stained using 0.1% Hoechst and counted on a haemocytometer using fluorescent microscopy.

#### Library Prep

Transposition reactions were performed using 300 thousand nuclei per reaction. 50 μL reactions of nuclei in transposition mix (25 μL 2x Tagmentation Buffer [Diagenode], 2.5 μL Tagmentase [Diagenode], 0.5 μL 10% Tween-20, 0.5 μL 1% Digitonin, 5 μL nuclease free H_2_O, 16.5 μL nuclei in PBTB) were incubated on a thermomixer at 1000 rpm and 37 °C for 30 minutes. Transposed DNA was purified using the DNA Clean & Concentrator-5 kit (Zymo Research) and used in subsequent PCR amplification. PCR was performed using Q5 High-Fidelity DNA Polymerase (New England Biolabs), Q5 Reaction Buffer (New England Biolabs), and Unique Dual Index PCR primers (Diagenode) under the following conditions: 72 °C adapter extension for 5 minutes and 98 °C denaturation for 30 seconds followed by 5 cycles of 98 °C denaturation for 10 seconds, 63 °C annealing for 30 seconds, and 72 °C extension for 1 minute. After 5 cycles, 5 μL of partially amplified libraries were used for qPCR reactions with the addition of 0.3 μL ROX and 0.1 μL 100x SYBR Green I, and brought up to 15 μL with water and supplemental PCR reagents to replicate original reagent concentrations. qPCR was performed for an additional 20 cycles, and the number cycles needed to reach around 1/3 saturation were used as additional cycles in the original PCR reaction. Amplified libraries were then size selected using SPRIselect beads at lower and upper concentrations of 0.55x and 1.8x to generate final libraries. Libraries were sequenced on a NovaSeq sequencer with a PE150 configuration for an expected yield of 11.11G per sample or roughly 36.7 million reads per sample.

#### Data Analysis

Reads were trimmed and filtered using Cutadapt and mapped to the Pmin_3.0 *Patiria miniata* genome (GCF_015706575.1) using bwa-mem2 (Vasimuddin et al., 2019). Mapped reads were filtered using Samtools (Danecek et al., 2021), and open chromatin regions were identified using MACS2 (Zhang et al., 2008). A set of consensus open chromatin regions were generated as the set present in at least two technical replicates of the same sample and used as input for differential accessibility analysis using the R package DiffBind (Rory Stark, 2017), which uses DESeq2 (Love et al., 2014). Transcription factor footprinting was performed using TOBIAS (Bentsen et al., 2020), and input position frequency matrices (PFMs) were the 2024 non-redundant vertebrate PFMs from JASPAR.

### In situ hybridization

#### DIG Probe Synthesis

cDNA was generated using RNA extracted from either 9 dpf or 3 dpb *Pmin* larvae with the Verso cDNA Synthesis kit (Thermo Scientific) according to the manufacturer’s protocol. Primers were then used to amplify regions of interest from cDNA. Each primer pair includes an SP6 promoter sequence added on to the end of reverse primers. Reverse transcription reactions were performed on amplified cDNA using the T7/SP6 DIG Labeling kit (Roche) according to the manufacturer’s protocol. Primers used for the construction of riboprobes, excluding SP6 promoter sequence, are as follows:

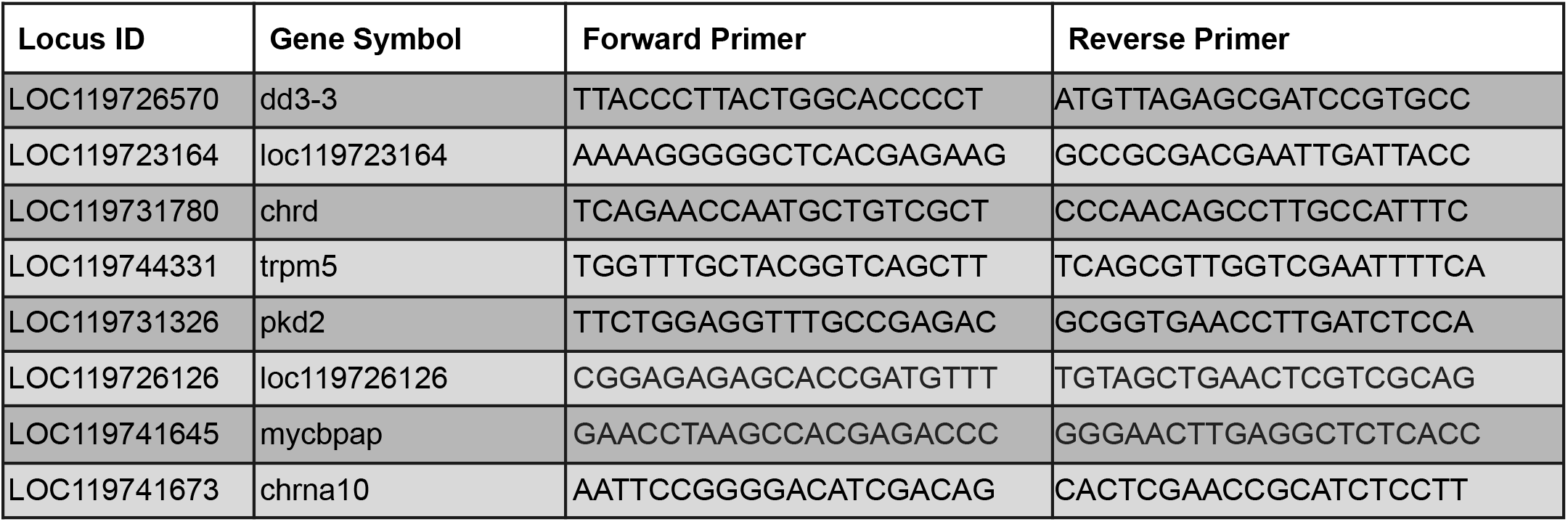
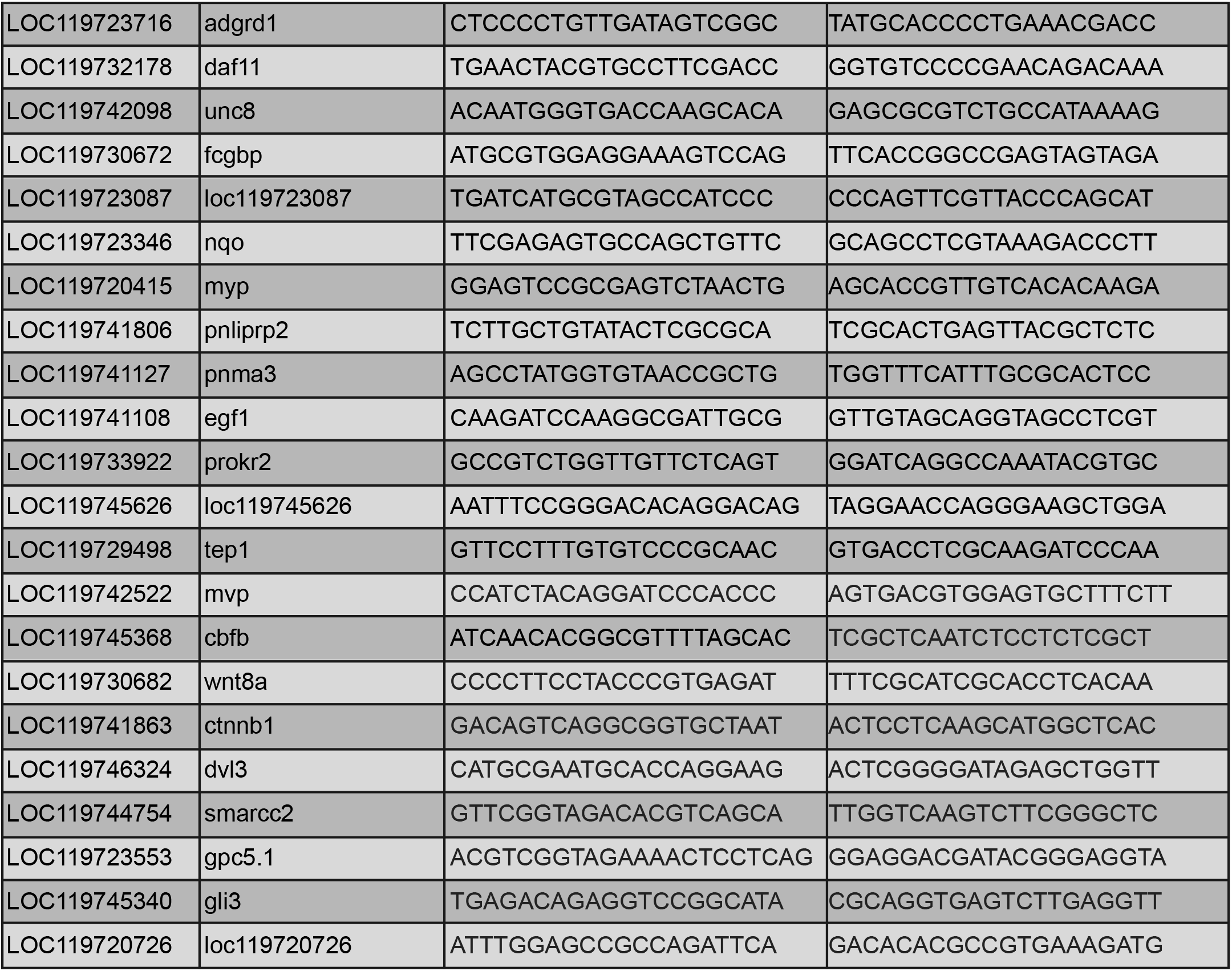

#### In situ hybridization

Fixed larvae were washed multiple times in 1x MOPS in situ buffer, followed by Hybridization buffer incubation overnight at 58 °C. DIG riboprobes were then added to a final concentration of 0.1 ng/μL, and samples were incubated at 58 °C for 3-5 days. Probes were washed out using 1x MAB in situ buffer, and samples were incubated in blocking buffer for 30 minutes at room temperature before incubation with 1:2000 dilution of anti-DIG AP FAB fragment in blocking buffer for 2 hours at room temperature. Antibody was washed out using 1x MAB in situ buffer, samples were washed with 1x AP buffer, and then samples were incubated in color reaction buffer (1x AP buffer, 50 mM MgCl_2_, 0.1% Tween-20, 10% Dimethylformamide, 0.35% Nitro Blue Tetrazolium, 0.35% 5-bromo-4-chloro-3-indolyl-phosphate) until localized signal was detected.

#### Reporter constructs

Reporter constructs were amplified from the *Pmin genomic* DNA using a nested PCR approach. Outer primers amplified regions from the genome, and nested primers with restriction sites on the ends amplified off outer PCR products. These products were then cloned into eGFP reporter constructs containing a minimal H2B promoter, SV40 PolyA sequence, ampicillin resistance, and a multiple cloning site. Inserts were digested with restriction enzymes, and empty vectors were digested with enzymes and also treated with calf intestinal phosphatase to prevent religation. Digested inserts and vectors were then combined with T4 ligase and incubated overnight at 4 °C. Ligase was subsequently deactivated at 65 °C for ten minutes, and ligation products were electroporated into electrocompetent DH10B *E. coli* and grown on ampicillin-containing plates. Transformed colonies were validated using PCR, and successful transformants were grown in liquid culture, miniprepped, and sequenced to verify successful insertion. Genomic coordinates of functionally validated regulatory elements and nested primers, without restriction sites, used in construct generation are as follows:

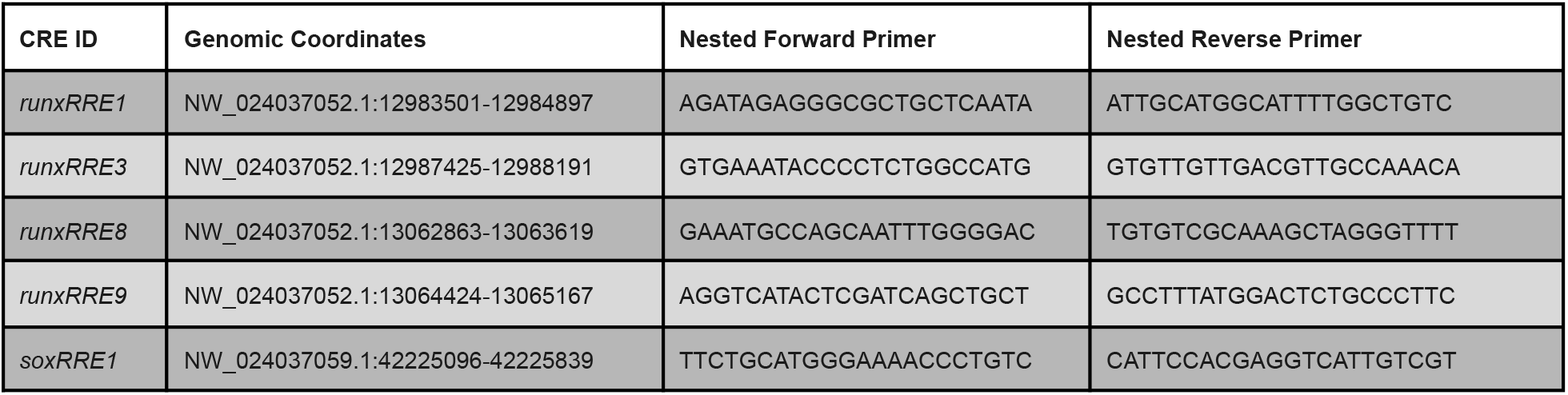

### Drug Treatment

To inhibit the MAPK/ERK signaling pathway, larvae were incubated with 15 μM U0126 1 hour prior to bisection. Larvae were then bisected in drug treated ASW and incubated overnight at 15 °C. 24 hours after initial treatment, 1 dpb larvae were fixed and used for in situ hybridization.

### Enhancer-Gene Association

To predict the genes regulated by enhancers, we identified the set of all transcription start sites (TSSs) both within 50 kb of an enhancer and within the boundaries established by the nearest up- and downstream intergenic spaces (Supp. Fig. S6). This allows for multiple predicted targets, in contrast to a nearest-TSS approach, but prioritizes genes based on proximity, yielding more reasonable sets of predictions in regions of dense gene population.

## Acknowledgements

I would like to express sincere gratitude to Dr. Anne Meyer for her assistance in establishing the data collection and preprocessing framework for the snRNA-seq experiments. I would also like to acknowledge Dr. Alyssa Lawler for her guidance and expertise in ATAC-seq data analysis. It is with the support of these and other contributors that this work was made possible.

---

**Supp Fig S1.**
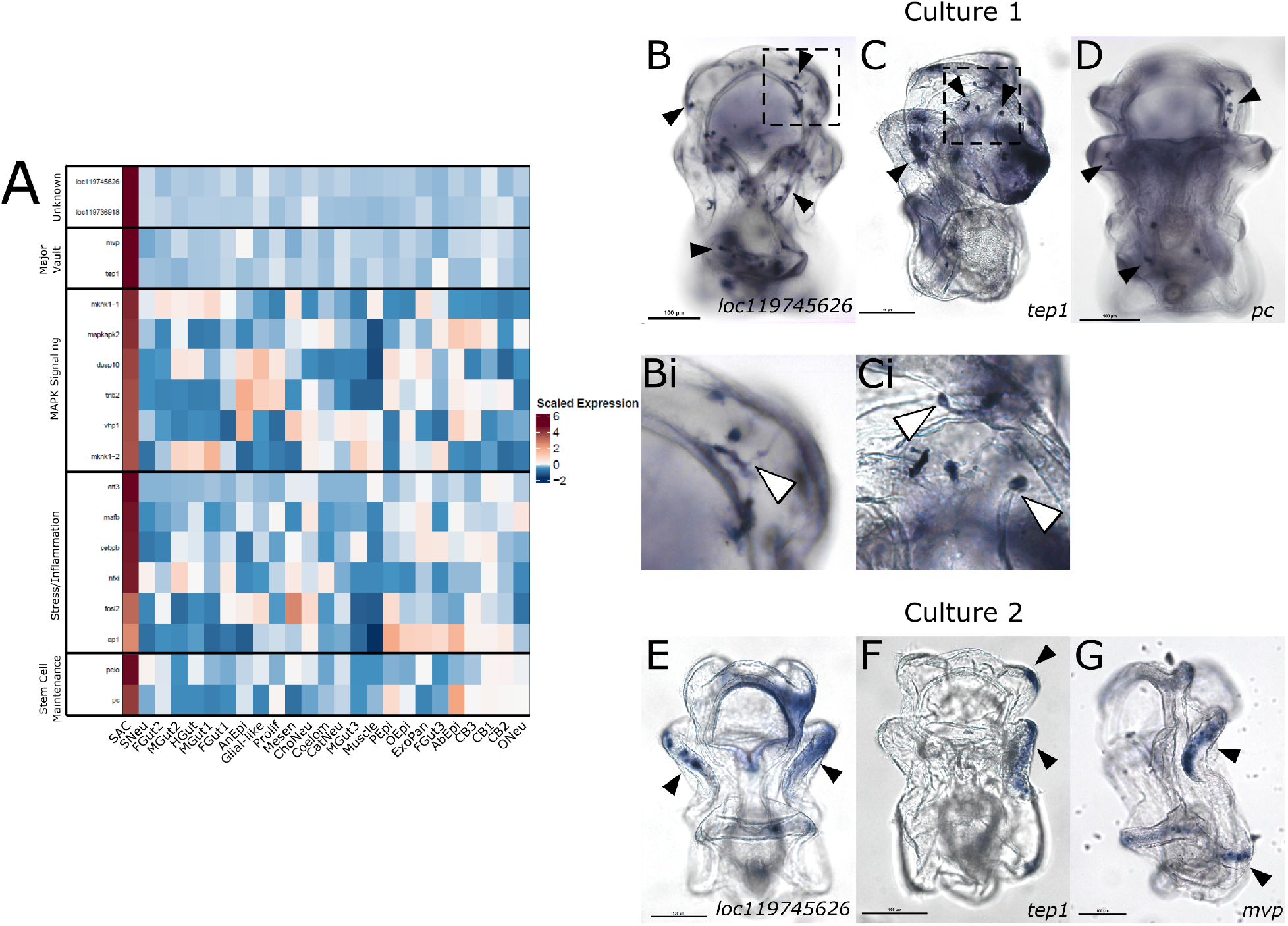
Stress-Associated Cell Markers and Variability. (A) Heatmap of SAC marker genes. Markers are along the y-axis organized by functional association. Clusters are along the x-axis. Color represents average expression in a cluster, scaled per row. Cluster name abbreviation conventions are similar to in Fig 1. (B-D) Marker localization using in situ hybridization within a single batch of fixed material for (B) loc119745626, (C) tep1, and (D) pc. Expression is seen as punctate cells throughout the body, indicated by black arrowheads. Membrane extensions (Bi) and teardrop morphologies (Ci) are observed as indicated with white arrowheads. (E-F) Marker localization using in situ hybridization within a different batch of fixed material for (E) loc119745626, (F) tep1, and (G) mvp. Expression is restricted to the ciliary bands and more diffuse.

**Supp Fig S2.**
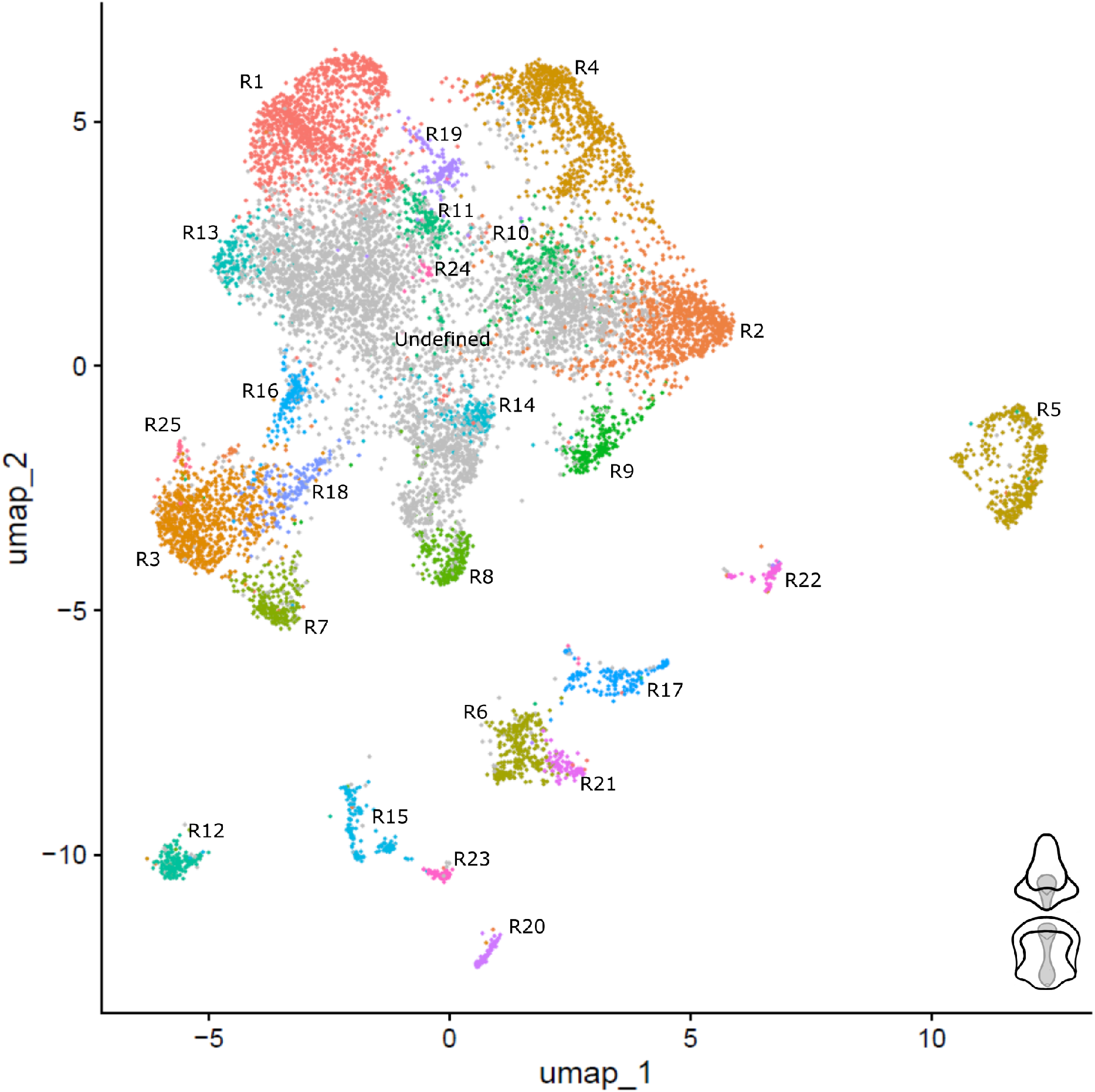
UMAP of 3 dpb Larval Nuclei. UMAP projection of nuclei from combined 3 dpb regenerating anterior and posterior fragments. 25 clusters are color coded and labelled in order of cluster size, with R1 being the largest. The undefined cluster not used in subsequent analyses is colored gray.

**Supp Fig S3.**
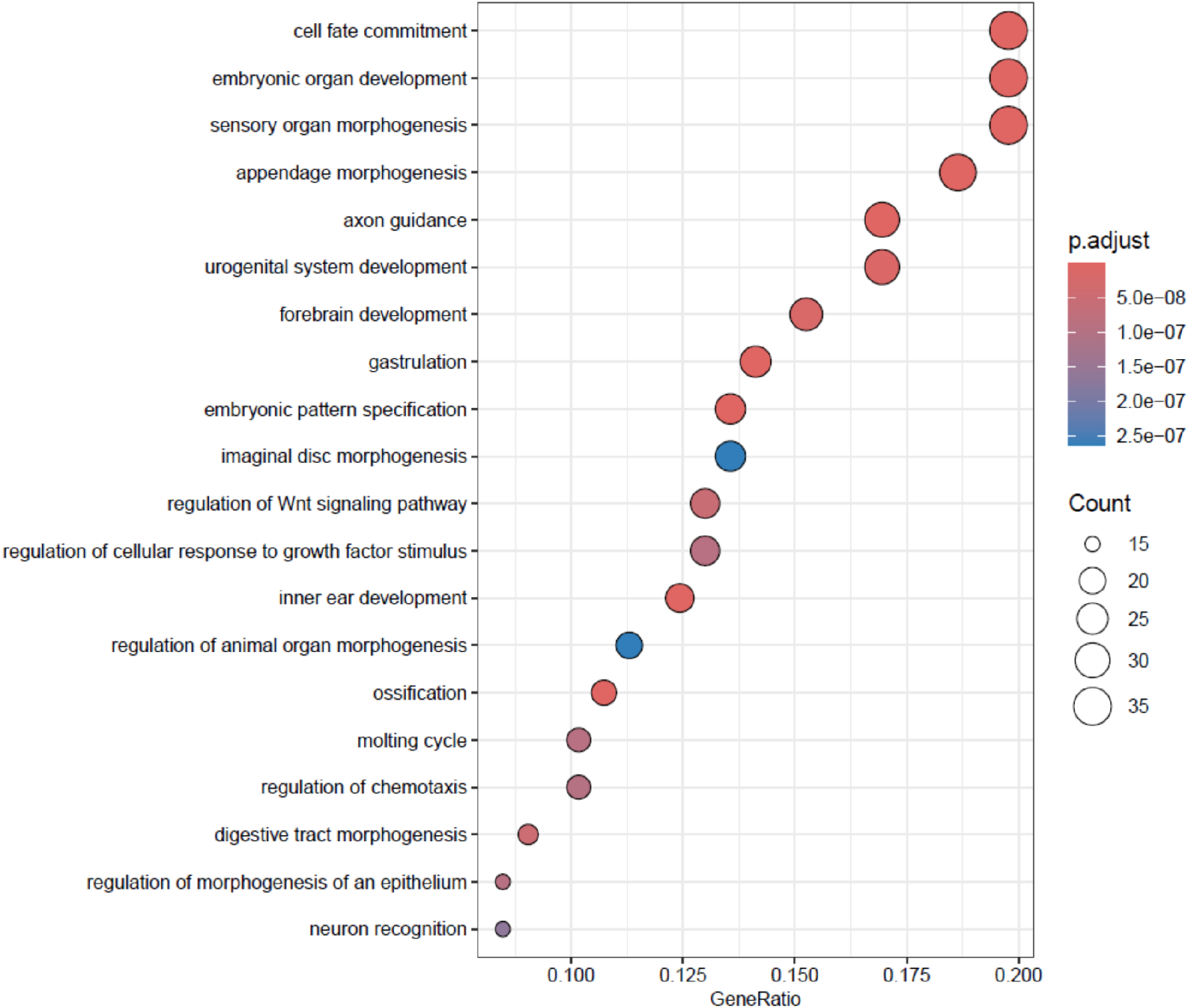
Enriched Functional Terms for R11. Dot plot for GO term enrichment for significant (adjust p < 0.05) R11 markers. Functional terms are on the y-axis, numbers of genes with a term is represented by dot size, ratio of gene number to total genes with that functional term is on the x-axis. Color represents significance.

**Supp Fig S4.**
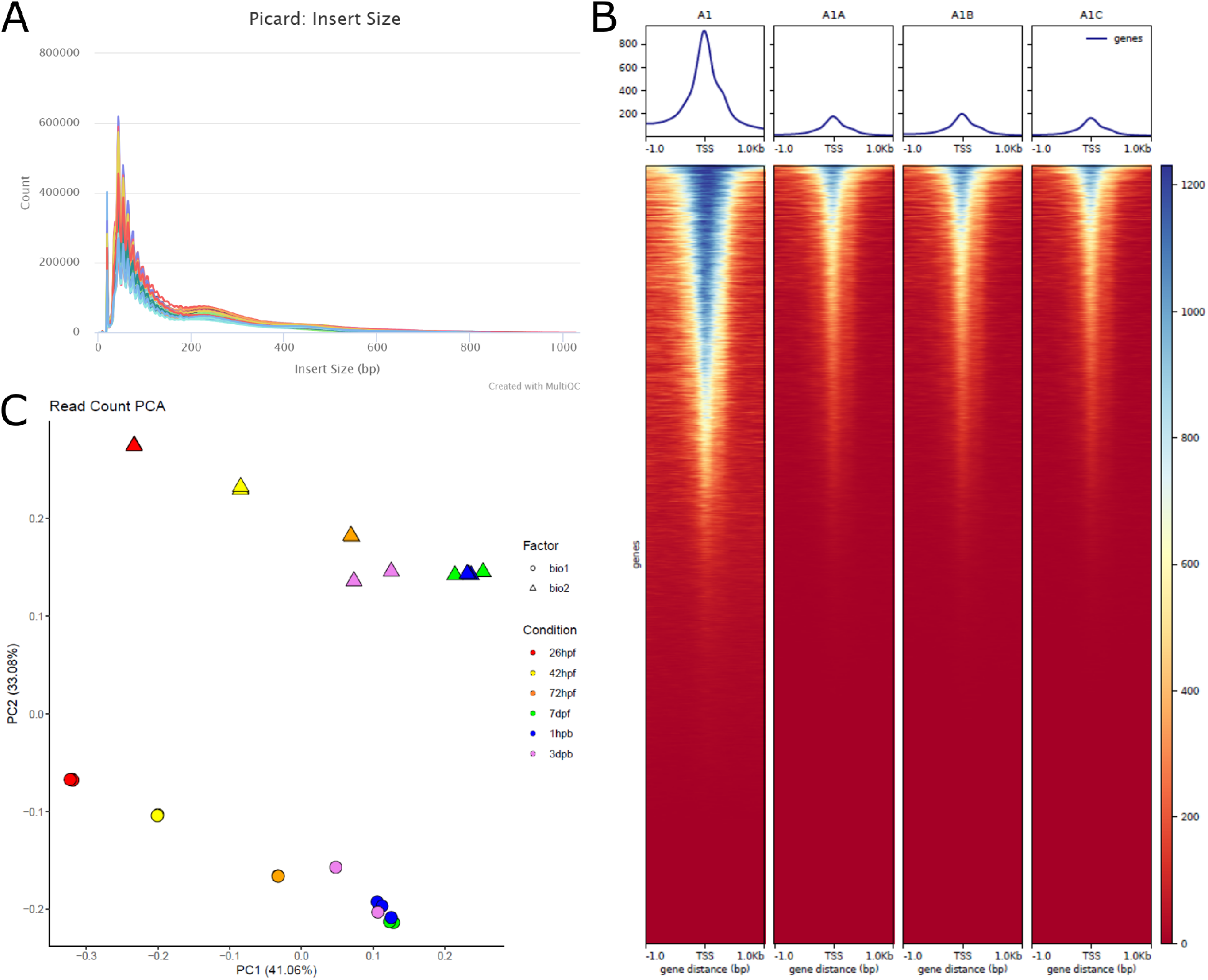
Quality Control for ATAC-seq Datasets. (A) ATAC-seq read insert size distribution. Trace color represents sample. Laddering is observed after the first peak, and subsequent peaks appear progressively weaker one nucleosome length apart (147 bp). (B) TSSE plot. Traces in square plots (top) represent average signal (y-axis) compared to relative distance to all transcription start sites (x-axis). Data for individual TSSs is depicted in long rectangular boxes (below). The x-axis is the same as for the average traces, the y-axis is made up of stacked lines that all represent different TSSs sorted from high to low signal, and color represents signal. Quality data will show increased signal at the TSS. (C) PCA plot of dimensionally reduced read data for different biological replicates (shape) and treatments (color). Biological replicates are quite different from each other, each of these follows a similar trajectory with the PCA space over time.

**Supp Fig S5.**
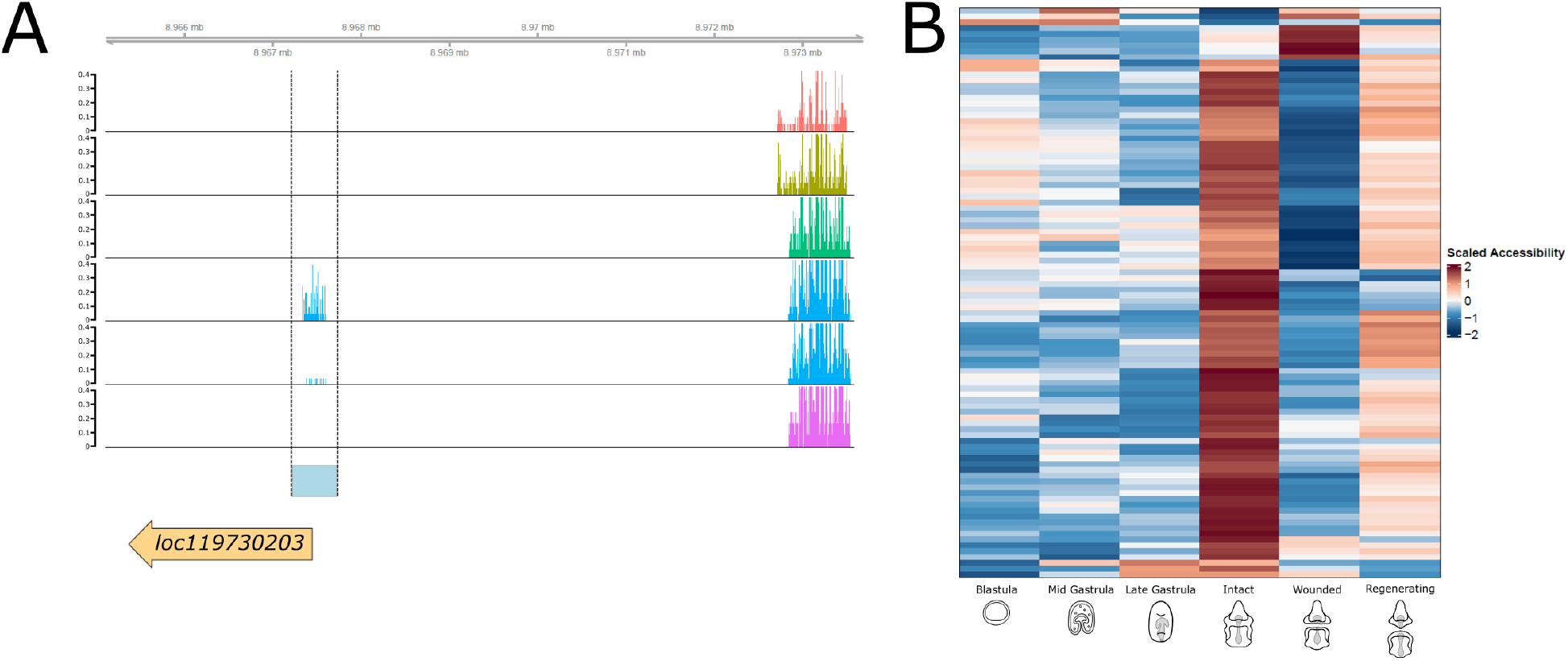
Wound Responsive Elements are Detectable by 1 hbp. (A) Genome track of ATAC signal around a representative OCR differentially accessible between intact and 1 hpb larvae. Samples are ordered along the y-axis in temporal order with blastula (red) on top, followed by mid gastrula (gold), late gastrula (green), intact larva (dark blue), 1 hpb larva (light blue), 3 dpb larva (purple). Signal value is indicated using scales on the left-hand side. The wound responsive element is indicated with vertical black lines. A nearby gene is present just 5’ of the wound responsive element. (B) Heatmap of the 98 OCRs differentially accessible between intact and 1 hpb wounded larvae. Samples are on the x-axis, OCRs on the y-axis clustered using Pearson correlation, and color represents accessibility, scaled per row. Most regions identified decrease in accessibility, and 9 increase.

**Supp Fig S6.**
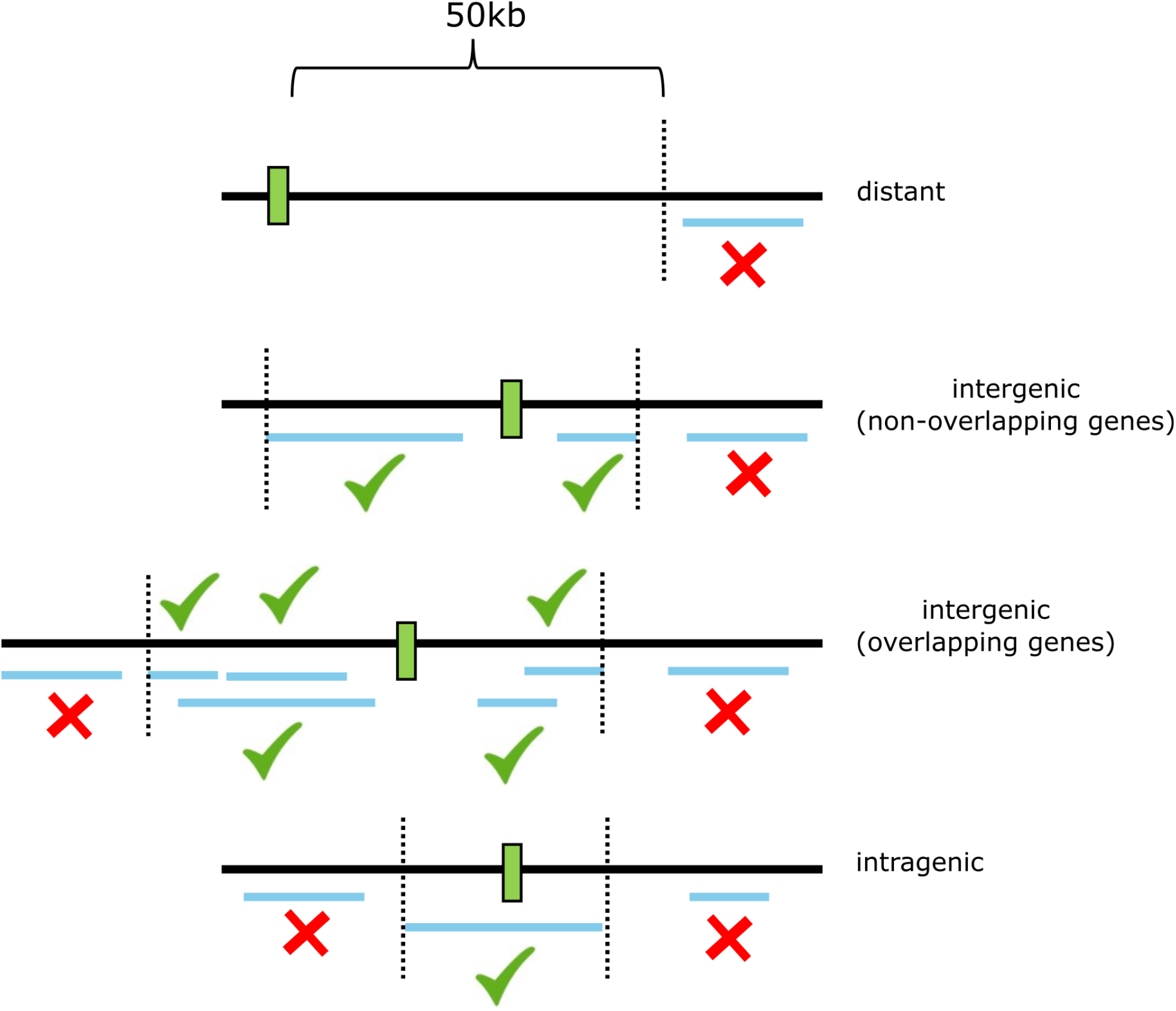
Regulatory Target Gene Prediction Approach. Schematic depicting approach for predicting regulatory target genes. OCRs (green box) lie within the genome (black line). Genes (blue lines) with TSSs within the bounds (dashed line) set by the next closest intergenic spaces or a hard cutoff of 50 kb are considered valid predictions.

**Supp Fig S7.**
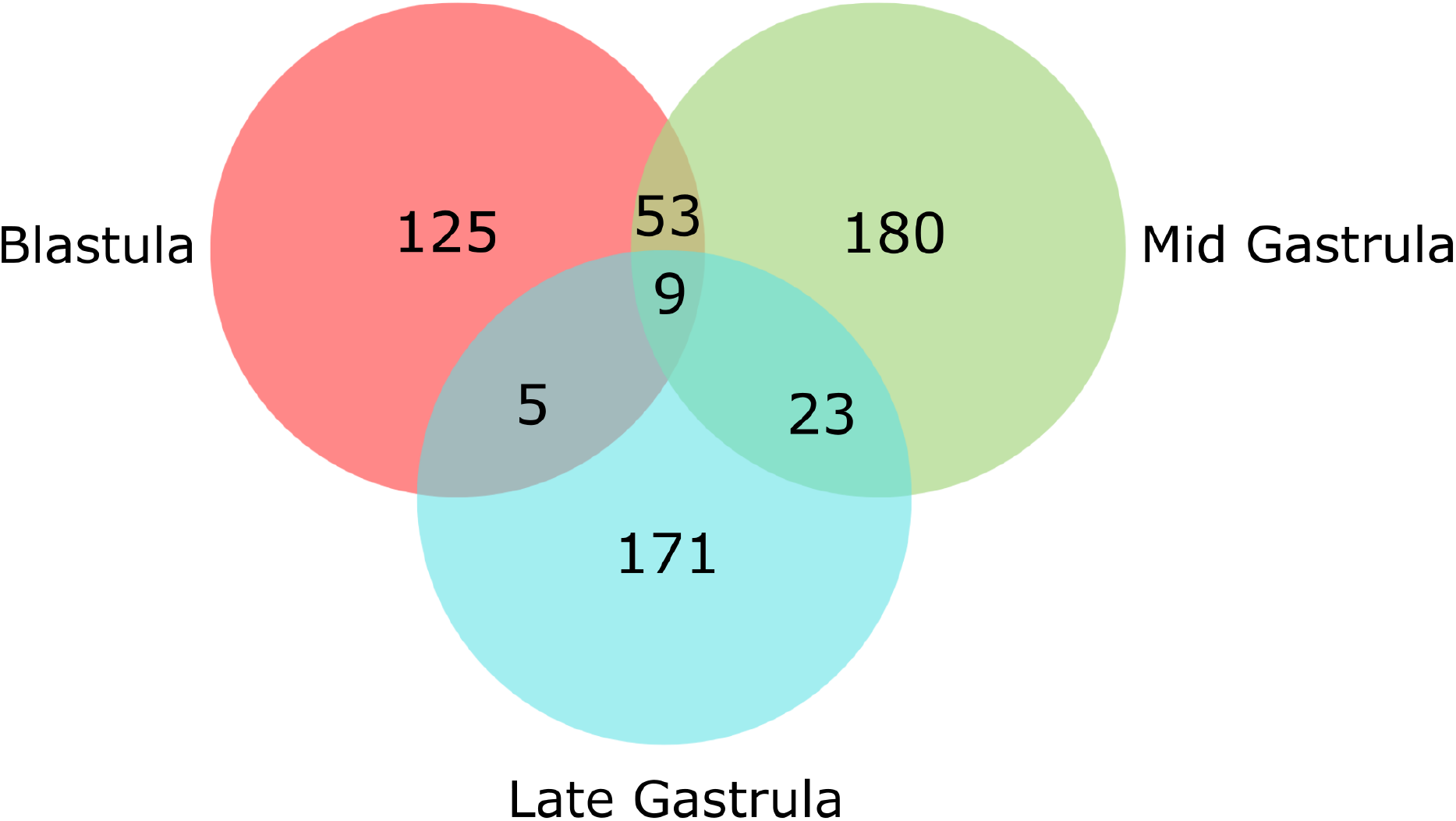
RREs Shared Between 3 dpb and Embryonic Time Points. Venn diagram of RREs in which 3 dpb accessibility is similar to (within 1.2-fold of) that of an embryonic sample. Most of these meet this criteria for only one embryonic stage.

**Supp Fig S8.**
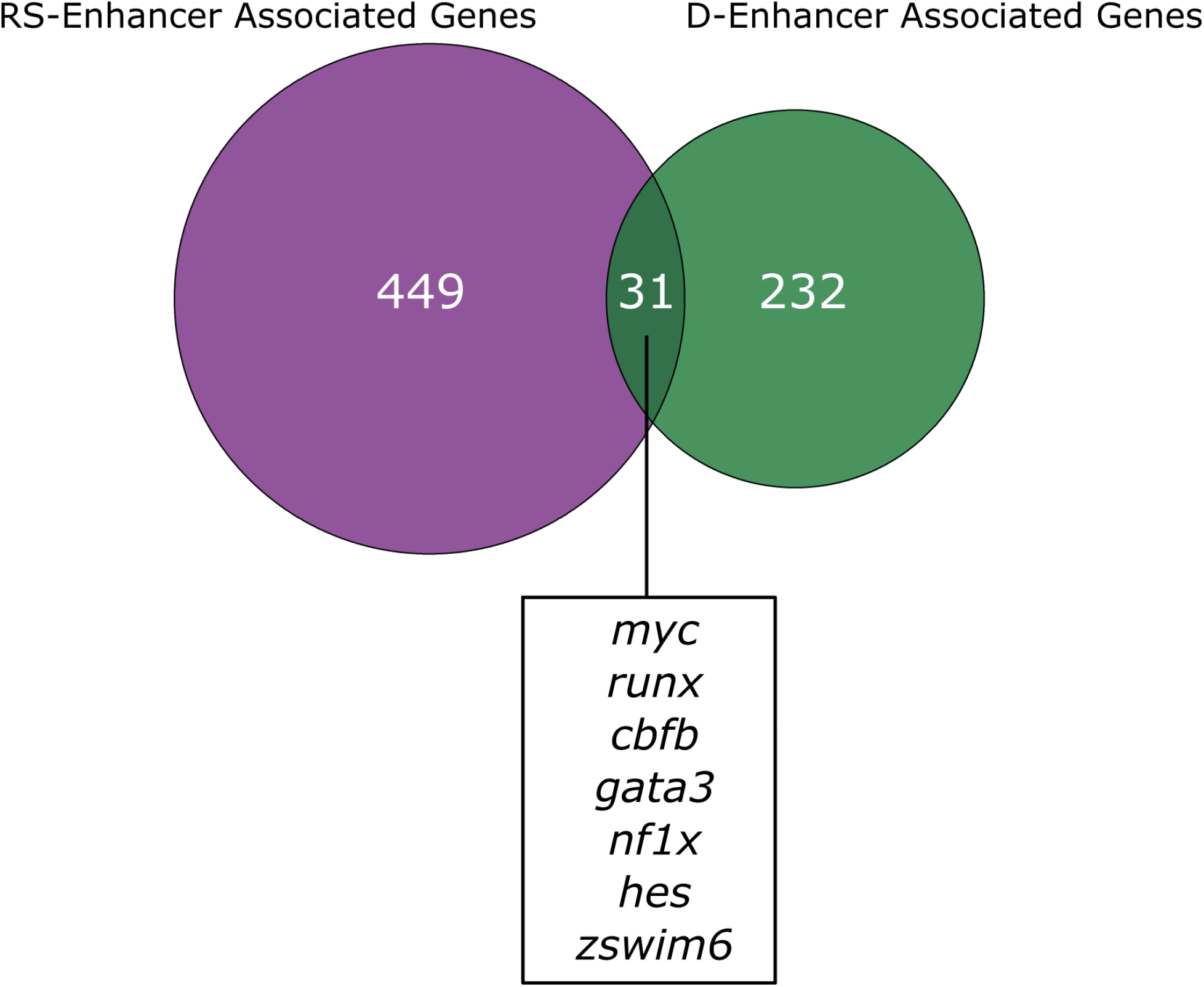
Overlap of Genes Predicted to be Regulated by RS- and D-Enhancers. Venn diagram comparing predicted regulatory gene targets of opening regeneration-specific RREs and opening developmental RREs. The vast majority of these genes are specific to either condition. 31 of these are regulatory targets of both RRE sets, which include runx, myc, cbfb, gata3, and others.

**Supp Fig S9.**
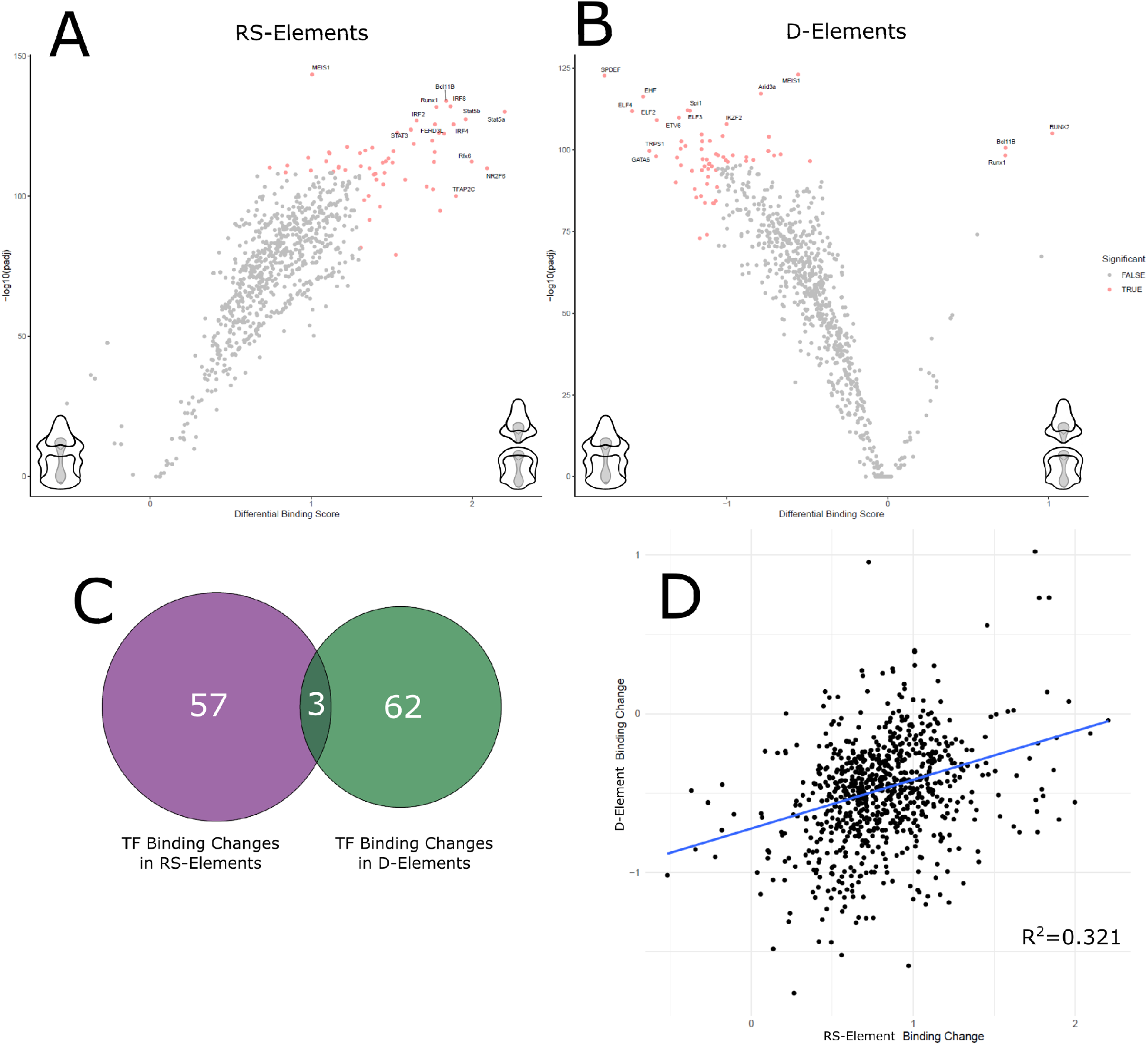
Differential Transcription Factor Usage Between RS- and D-Elements. (A-B) Volcano plots showing average change binding (x-axis) between intact and 3 dpb larvae for motifs within RS-elements (A) or D-elements (B). Most changes, and all significant changes (−log10(adjusted p) > 95th percentile, absolute differential binding score > 95th percentile), for RS-elements are increases in binding. The opposite is true for D-elements with the exceptions of Runx1, Runx2, and Bcl11b. (C) Venn diagram comparing the significant hits for A and B. Motifs that appear as significant increases or significant decreases in both conditions lie in the overlap, including 3 transcription factor motifs. Most other motifs do not have similar patterns of change between conditions. (D) Dot plot representing the differential binding scores for each motif, as in A and B, within RS-elements (x-axis) versus D-elements (y-axis). There is weak, yet positive, correlation (R2 = 0.321).

## Supplementary Text: single nucleus Atlas of 9 day binpinarria larva

### Non-Neural Ectoderm

Within the non-neural ectoderm, we detect three ciliary band clusters. All of these show high expression of *forkhead box J1* (*foxj1*) (Fig. 1B), a known marker of the larval ciliary bands (Yankura et al., 2010). Cells of all ciliary band clusters are found throughout both the pre- and post-oral ciliary bands, generally with a decrease in marker expression closer to the anterior pole (Fig. 1G-I).

Interestingly, ciliary band 1 shows strong and highly specific expression for many c-type and lactose-binding lectins, which are associated with innate immunity in echinoderms (L. C. Smith et al., 2010).

We also detect four clusters representing different spatial domains of the non-ciliated epidermis. Cells of the anterior epidermis express *protein DD3-3* (*dd3-3*) (Fig. 1C) as well as *frizzled class receptor 5* (*fzd5*) (Fig. 1B), a known anterior marker for both embryonic and regenerating *P. min* (Cary et al., 2019; McCauley et al., 2013). The posterior epidermis expresses *loc119723164* (Fig. 1D). We also detect *Wnt family member 3* (*wnt3*)*, wnt10b,* and *homeodomain protein* (*Hbox7*), which are expressed in posterior tissues of *P. min* embryos and sea urchin larvae (Annunziata & Arnone, 2014; Cui et al., 2017; McCauley et al., 2013; J. Smith et al., 2008; Yamazaki et al., 2012). The oral ectoderm includes the domain ventral to and enclosed by the post-oral ciliary band, with the exception of the most anterior tissues (Fig. 1E), and expresses *chordin* (*chrd*), which is expressed dorsally in vertebrates but ventrally in flies and other echinoderms (Meinhardt, 2015). Conversely, the aboral ectoderm, marked by *transient receptor potential cation channel subfamily M member 5* (*trpm5*), includes the domain dorsal to and enclosed by the post-oral ciliary band (Fig. 1F). Interestingly, marker expression for both the oral and aboral ectoderm is detected within the domain enclosed by the pre-oral ciliary band (Fig. 1E-F).

### Neural Ectoderm

We detect 5 neural clusters in our dataset. Marker genes for all clusters each include one or more homologs of both *elav* and *synaptotagmin* (*syt*), which has been used to identify neurons in larval echinoderms including *P. min* (M. Zheng et al., 2022), and between 5 and 12 voltage-gated or -dependent ion channel genes, supporting our annotations (Supp. Data S1). However, it is important to note that homologs for *elav* and *syt* appear as significant markers (adjusted *p* < 0.01, Wilcoxon rank-sum test) for other non-neural clusters including midgut clusters 1 and 3, the hindgut, exocrine pancreas-like cells, mesenchyme, and muscle (Supp. Data S1). Cholinergic neurons, expressing *choline O-acetyltransferase* (*chat*), are located along the ciliary bands, particularly the medial loop of the post-oral ciliary band and the transverse segments of both ciliary bands (Fig. 1N), and catecholaminergic neurons, expressing *tyrosine hydroxylase* (*th*), have a similar spatial pattern with additional localization to the lower lip of the mouth and hindgut opening (Fig. 1O), both in agreement with recent neuronal characterizations in *P. min* (Pagowski, 2024).

Serotonergic neurons, expressing *tryptophan 5-hydroxylase 1-like* (tph1), are located in the dorsal ganglia along the anterolateral post-oral ciliary band (Fig. 1P). This agrees with earlier characterization of this cell type in *P. min* (M. Zheng et al., 2022).

We also identified a cluster of neurons located along the entire border of the mouth, which we annotate as oral neurons (Fig. 1Q). GO analysis shows an enrichment of genes predicted to be involved in ion homeostasis and chemical sensory perception (Supp. Data S2). Another neural-like cluster strongly expresses genes involved in cholesterol biosynthesis according to GO analysis (Supp. Data S2), which we annotate as glial-like, as this is a common feature of glial cells (Vance et al., 2005). Cells of this cluster, which is the largest of the neural ectoderm, are found throughout the ciliary band and exhibit highly specific expression of *glial cells missing transcription factor-like* (*gcml*) (Fig. 1B, 1R), a major marker of immune pigment cells in sea urchins and of all non-serotonergic neuronal types during *P. min* embryogenesis (Meyer et al., 2023).

### Endoderm

We detect 8 distinct endodermal clusters in our snRNA data, including all 3 compartments of the tripartite gut–the foregut, midgut, and hindut–as well as an exocrine pancreas-like population. The foregut is divided into 3 clusters defined by highly specific marker genes (Fig. 1B). The foregut 1 marker, *IgGFc-binding protein* (*fcgbp*), clearly delineates the tissue (Fig. 1S). Spatial expression was not validated for foregut clusters 2 and 3. However, foregut 2 expresses *AT-rich interactive domain-containing protein 3A* (*arid3a*), which marks the foregut and right coelom during *P. min* embryogenesis (Meyer et al., 2023), supporting our annotation.

The midgut is also split into 3 clusters that spatially overlap (Fig. 1T-V).

Midgut 1, the largest of the three, exhibits the highest expression of *GATA binding protein 6* (*gata6*) and *RNA-binding protein Nova-2-like* (*nova2*) (Fig. 1B), which are known midgut markers in sea star development and larval urchins (Meyer et al., 2023; Röttinger et al., 2006). Markers for midgut 1, however, are not notably specific, and are highly expressed in the other midgut clusters (Fig. 1B). Midgut 2, which has the most specific gene markers among the midgut clusters (Fig. 1B), is enriched for GO terms associated with molecular transport and the brush border, suggesting potential nutrient absorptive function (Supp. Data S2). Midgut 3 expresses *quinone oxidoreductase-like* (*nqo*), which is involved in detoxification (Ross & Siegel, 2021).

The hindgut expresses *major yolk protein* (*myp*) (Fig. 1W), which agrees with previous expression data in *P. min* (Zazueta-Novoa et al., 2016). *Pancreatic and duodenal homeobox 1-like* (*pdx1l*) and *caudal type homeobox 1-like* (*cdx1l*), which are both expressed in the hindgut of larval *S*. *purpuratus* (Paganos et al., 2021), are also markers for this cluster (Fig. 1B, Supp. Data S1). We also detect an exocrine pancreas-like cell type, which has been described in *S. purpuratus*, the first example in echinoderms (Perillo et al., 2016). Similar to in urchins, these cells are located at the anterior portion of the midgut (Fig. 1X) and express various proteases, some of which are known to be expressed in this population, including *carboxypeptidase B-like* (*cpb*) and *aqualysin-1-like* (*pstl1*); as well as lipases, including *pancreatic lipase-related protein 2-like* (*pnliprp2*) (Fig. 1B) (Paganos et al., 2021; Perillo et al., 2016).

### Mesoderm

Within the mesoderm, we detect the coelom, mesenchyme, and muscle.

The coelomic marker, *paraneoplastic antigen Ma3 homolog* (*pnma3*), is localized to the entire length of both bilateral coelomic pouches as well as the posterior coelom (Fig. 1J). Within the marker gene set, we also detect transcription factors previously confirmed in the coeloms of *P. min* and/or other larval and embryonic echinoderms including *alx1*, *homoeobox A9* (*hoxa9*), *GLI family zinc finger 3* (*gli3*), *forkhead box F1* (*foxf1*), *SIX homeobox 1* (*six1*), *paired box 6* (*pax6*), and *transcription factor AP-2 alpha* (*tfap2a*) (Supp. Data S1) (Arenas-Mena et al., 2000; Cary et al., 2019; Hara et al., 2006; Koop et al., 2017; Luo & Su, 2012; McCauley et al., 2012; Meyer et al., 2023; Tu et al., 2006; Yankura et al., 2010).

The mesenchyme is distributed throughout the blastocoelar cavity and in the coelomic epithelium (Fig. 1K). Markers for this cluster have been shown to have similar distribution in *P. min* larvae or to be expressed in immune-mesenchyme populations in embryonic *P. min* snRNA-seq data, as with *ets1*, *erg*, *GATA binding protein 3* (*gata3*), *TEK receptor tyrosine kinase* (*tek*), and *dual oxidase 1* (*duox1*) (Cary et al., 2019, Meyer et al., 2023), and several markers have also been shown or predicted to be important in immune cell development and function in other echinoderms, including *gata3*, *interferon regulatory factor 4* (*irf4*), and *tek*, as well as scavenger receptor cysteine-rich-domain, von Willebrand factor domain, and sushi domain containing proteins (Supp. Data S1) (Chiaramonte et al., 2019; Furukawa et al., 2012; Huang et al., 2010; Pancer et al., 1999; L. C. Smith et al., 2018). The muscle is located in two longitudinal bands on the dorsal side of the animal as well the posterior portion of the foregut, likely allowing the larva to swallow and to bend at the midline (Fig. 1L).

### Proliferating Cells

We also detect the presence of proliferating cells, which express genes classically associated with mitosis such as *inner centromere protein* (*incenp*), *cyclin B2* (*ccnb2*), and *aurora kinase A* (*aurka*) (Fig. 1B). Because these were captured using nuclear extraction, we expect these to be in prophase or earlier before the breakdown of the nuclear envelope. Localization of these cells through in situ hybridization was unsuccessful, but we expect these cells to be located throughout the body, particularly the ectoderm, at this stage (Cary et al., 2019).

## Notes

### Competing Interest Statement

The authors have declared no competing interest.

